# Cell Type-specific Genome Scans of DNA Methylation Diversity Indicate an Important Role for Transposable Elements

**DOI:** 10.1101/801233

**Authors:** Önder Kartal, Marc W Schmid, Ueli Grossniklaus

## Abstract

The epigenome modulates the activity of genes and supports the stability of the genome. The epigenome can also contain phenotypically relevant, heritable marks that may vary at the organismic and population level. Such non-genetic standing variation may be relevant to ecological and evolutionary processes. To identify loci susceptible to selection, it is common to profile large populations at the genome scale, yet methods to perform such scans for epigenetic diversity are barely tapped. Here, we develop a scalable, information-theoretic approach to assess epigenome diversity based on Jensen-Shannon divergence (JSD) and demonstrate its practicality by measuring cell type-specific methylation diversity in the model plant *Arabidopsis thaliana*. DNA methylation diversity tends to be increased in the CG as compared to the non-CG (CHG and CHH) sequence context but the tissue or cell type has an impact on diversity at non-CG sites. Less accessible, more heterochromatic states of chromatin exhibit increased diversity. Genes tend to carry more single-methylation polymorphisms when they harbor gene body-like chromatin signatures and flank transposable elements (TEs). In conclusion, the analysis of DNA methylation with JSD in *Arabidopsis* demonstrates that the genomic location of a gene dominates its methylation diversity, in particular the proximity to TEs which are increasingly viewed as drivers of evolution. Importantly, the JSD-based approach we implemented here is applicable to any population-level epigenomic data set to analyze variation in epigenetic marks among individuals, tissues, or cells of any organism, including the epigenetic heterogeneity of cells in healthy or diseased organs such as tumors.

## Introduction

The ongoing development of sequencing-based functional genomics has a tremendous impact on the study of gene regulation. Nowadays, we can get an almost comprehensive, genome-wide readout of gene expression and chromatin states. This technological progress not only produces genomic data sets for more and more organisms but enables us to profile gene regulation at the level of organs, tissues, and even individual cells. However, new technologies beget new problems. We are confronted with multidimensional data sets and a sophisticated sampling situation that involves population structure, cell heterogeneity, temporal change, and technical bias, raising new questions about how regulatory factors vary within and across these different levels. The issue of diversity and its apportionment has a prominent place in population genetics. To find loci under selection, genomes are scanned for conserved or polymorphic sites using metrics for genetic differentiation like *F*_ST_ [1], which takes into account the association of alleles with environmental factors and population structure. The translation of these population-level approaches to genome-wide chromatin marks is still missing but the data currently available provides a solid foundation for developing and testing measures of epigenetic diversity.

DNA methylation has been studied extensively in humans and several model organisms at genome scale. Whole-genome sequencing of bisulfite-converted DNA (BS-seq) [2, 3, 4] provides accurate, genome-wide maps of the chemically modified cytosine base 5-methylcytosine (5mC) at single-base resolution, so-called methylomes. Although 5mC is not ubiquitous, it is widespread in higher eukaryotes [5]. In vertebrates, it is mainly found at CpG dinucleotides (CG context), whereas plants harbor 5mC also in the CHG and CHH context (H being A, C, or T).

DNA methylation can interfere with gene expression and contributes to the silencing of repetitive elements [6]. Moreover, the methylation landscape must be actively controlled throughout the life cycle by an enzymatic machinery. In mammals, extensive reprogramming takes place during primordial germ cell development and early embryogenesis [7] and the methylome also changes during tumorigenesis [8]. In plants, reprogramming is less extensive and the details are yet unclear but some epigenetic marks get reprogrammed during reproduction [9, 10, 11]. A drastic perturbation of the methylation pathways is lethal to mammalian embryos [12] and can lead to sterility and developmental aberrations after inbreeding in plants [13]. These severe effects illustrate an essential role for DNA methylation, not only for the activity of specific genes but for the integrity of chromatin as a whole.

Due to its correlation with fitness-relevant traits and its susceptibility to stress, DNA methylation has attracted considerable interest in evolutionary biology as a mediator of soft inheritance. For evolutionary studies, *Arabidopsis thaliana* is an excellent model for organisms with a full-featured methylation machinery. It has a small genome, a short life cycle, and large populations that harbor methylation polymorphisms of natural [14, 15, 16, 17] as well as artificial origin [18, 19] are available.

To enable genome scans of methylation diversity at single-base resolution, we use a non-parametric approach based on Jensen-Shannon divergence (JSD), a divergence measure in information theory with unique properties [21, 22]. JSD measures the loss of information (or, equivalently, the increase in uncertainty) if a set of distinct, information-carrying units is pooled. To define JSD formally, the concepts of probability distribution and Shannon entropy [23] are necessary (see Methods for the details) but Figure 1a illustrates the calculation of JSD geometrically. In this example, the three probability distribution functions (PDFs) are a sample from the population whose diversity is estimated by measuring the length of the dashed blue line. In the case of methylome data, the PDF of each individual is derived from the count data in a methylation table. A methylation table assigns two numbers to each cytosine site in the reference genome, the count of methylated and unmethylated reads. Therefore, in a population sample with *s* methylomes, each site is associated with a contingency table of 2×*s* entries. JSD is used to map this site-specific table to a site-specific number that reflects whether the methylation state at the given site is conserved (dip in JSD) or diversified (peak in JSD). Table 1 exemplifies the computation of JSD at a single site using the plug-in, or empirical, estimator (see Methods).

**Table 1:** How to compute the terms of JSD at site *i* for a sample of three methylomes. The result is 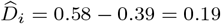. NA: not applicable.

| | $n_1$ | $n_2$ | $\mu$ | $\pi$ | $H$ | $\pi H$ |
| --- | --- | --- | --- | --- | --- | --- |
| $P_{i1}$ | 15 | 0 | 1.00 | 0.42 | 0.00 | 0.00 |
| $P_{i2}$ | 11 | 1 | 0.92 | 0.33 | 0.41 | 0.14 |
| $P_{i3}$ | 5 | 4 | 0.56 | 0.25 | 0.99 | 0.25 |
| $\langle P_i \rangle$ | 31 | 5 | 0.86 | NA | 0.58 | 0.39 |

**Figure 1.**
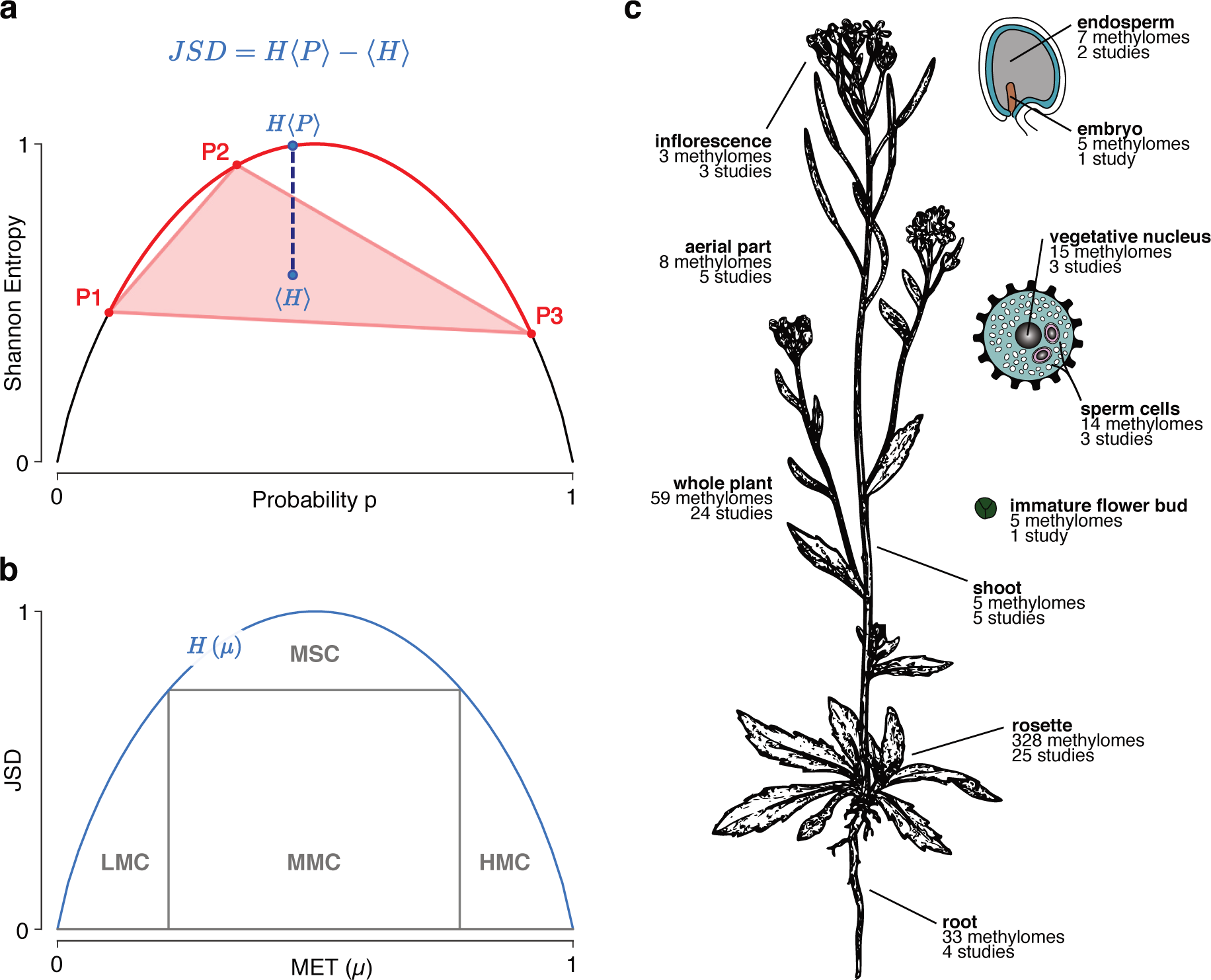
Application of Jensen-Shannon divergence to DNA methylation. **a** Geometric explanation of Jensen-Shannon divergence (JSD) in terms of Shannon entropy. The graph shows the entropy of a binary distribution (with probabilities *p* and *q* = 1 − *p*) in terms of *p*. For the purpose of illustration, the red dots designate the coordinates of three distributions, *P*_1_; *P*_2_; *P*_3_ that define a red curve segment and a corresponding polygon (red triangle). The curve segment constrains the entropy of the mixture distribution, *H*〈*P*〉, while the polygon constrains the corresponding average entropy, 〈*H*〉. The exact location of both JSD terms depends on the weights. For equally weighted distributions (here *π*_*i*_ = 1/3 for all *i*), their locations are given by the blue dots. The corresponding distance (length of the dashed line) equals JSD. **b** Phase plane in terms of weighted average methylation MET (*μ*) and diversity index JSD. The methylation state of a cytosine in the population is represented by a point at or below the graph. Four regions of interest are highlighted: three regions with JSD below ≈ 0:7, LMC (low-methylated cytosines), MMC (medium-methylated cytosines), and HMC (high-methylated cytosines); a region with high JSD for MSCs (metastable cytosines). **c** Overview of methylome data sources. 482 Arabidopsis methylomes from 75 different studies have been analyzed. All methylomes derive from wild type plants of the Columbia 0 (Col-0) ecotype. The image is modified based on [20].

We have analyzed methylation diversity in different parts of *Arabidopsis* as summarized in Figure 1c. We distinguish the methylomes according to the source of the corresponding DNA, that is according to which cell type, tissue, or organ the DNA was extracted from. Our analysis emphasizes the role of sequence context, chromatin accessibility, and genomic location, particularly the proximity to transposable elements (TEs), in shaping DNA methylation diversity.

## Results

### Genomic spectrum of methylation diversity

This section summarizes the genome-wide features of methylation diversity depending on genomic source and sequence (C) context. The source is expected to affect methylation diversity because the activity of genes differs between tissues and cell types. The C context is expected to affect diversity because different mechanisms are responsible for maintaining methylation in the CG, CHG, and CHH context [24].

To get a birds-eye view of the methylation state in a population, we base our analysis on the phase plane spanned by methylation bias and diversity, as measured by MET and JSD (see Methods). As indicated in Figure 1b, the phase plane allows classifying C-sites into different cytosine types (C types). C-sites with low to moderate diversity are divided according to low (LMCs), medium (MMCs), or high (HMCs) methylation bias; C-sites with exceptionally high JSD are classified as metastable Cs (MSCs) that segregate across the population (see Methods). We quantified the proportion of C types both for the whole genome and for non-overlapping, genomic intervals to analyze their enrichment in specific regions.

#### Diversity is higher in the CG than the non-CG context

Figure 2a shows the rosette leaf phase plane for each context. It illustrates what we find in all analyzed sources, namely that the methylome is stable over a wide range of MET values. Both, the histograms in the phase plane margins of Figure 2a and the empirical cumulative distribution functions (Fig. 2b) illustrate that more than 90% of C-sites have a JSD below 0.2 bit in each context. The low proportion of MSCs (see Fig. 2c) underlines that the population shows only weak to moderate segregation at the majority of sites, irrespective of C context. The C-sites with low to moderate JSD are mainly LMCs, as expected for the largely unmethylated genome of *Arabidopsis*. The proportion of LMCs is highest in the CHH and lowest in the CG context. The LMCs are also responsible for the positive correlation between MET and JSD (spearmanr in Fig. 2a). The correlation is consistently positive for LMCs regardless of source and context; the correlation varies by source and context, however, in other subregions of the phase plane (Fig. S1): for HMCs, MET and JSD are rather negatively correlated; for MMCs, MET and JSD are virtually uncorrelated; for MSCs, the correlation is weakly (CG context) or moderately positive (non-CG context).

**Figure 2.**
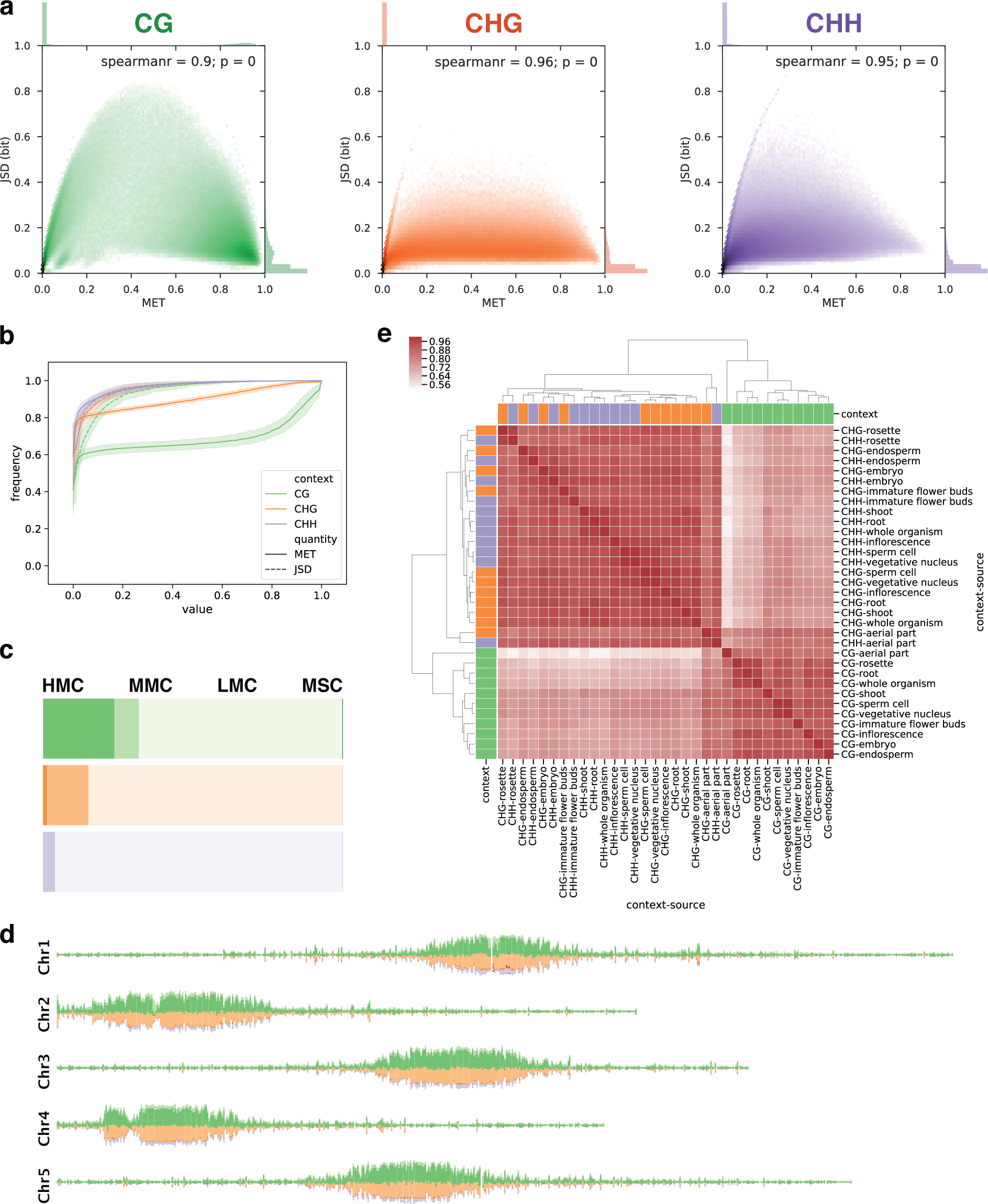
Genomic spectrum of methlation diversity. **a** Methylome phase plane for rosette leaves. The margins display the distribution of MET and JSD, respectively. SciPy (scipy.stats.spearmanr) was used to compute the Spearman correlation coefficient and *p*-value. **b** Empirical cumulative distribution function for MET and JSD in all three C contexts (color-coded). The line and error band display the mean and standard deviation of the data of all sources. **c** Proportions of C types for rosette leaves based on phase plane partitioning. **d** Rosette chromosome tracks for the proportions of C types at non-overlapping, 50 kb intervals over all nuclear chromosomes (1 to 5 from top to bottom). Bars for LMCs are omitted, that is smaller bars indicate a high proportion of LMCs. **e** Hierarchical clustering of genomic JSD signal for all source-context combinations using the Spearman correlation over non-overlapping, 50 kb intervals.

As shown in Figure 2b, the cumulative distribution functions of MET and JSD, respectively, do not differ by C context up to the median value of the distribution (specified at frequency= 0.5). However, there is a tendency for higher JSD in the CG context indicated by the long tails of the distributions at higher percentiles, respectively. The Mann-Whitney *U* test has confirmed that there is strong evidence for higher JSD in the CG context (see Table S2). The propensity for higher MET in the CG context is also reflected by the different proportions of C types: around 24% of the CG sites are HMCs, as opposed to only around 1% of the CHG and only 0.02% of the CHH sites (see Fig. 2c and Table S3 for a comprehensive summary). In fact, the joint distribution in the rosette phase plane for CGs is bimodal with one peak at LMCs and another one with slightly higher JSD formed by HMCs.

Note that the 328 rosette methylomes considered here originate from leaves that have been harvested at different developmental stages (9), from plants that have been grown in different labs (13), under different stresses (7), and photoperiods (3). The stability of the methylome despite this heterogeneity suggests that environmental conditions have a minor effect on methylation at large—genotype and cell-type specific regulation play the main role in shaping the methylation landscape.

#### Non-CG sites segregate within rather than among individuals

The rosette phase plane demonstrates another striking difference between the CG and CHG (and, to some extent, CHH) context around MET= 0.5; non-CG sites mainly occupy regions of lower JSD. First of all, a low JSD for a given site means that there is weak segregation across the population. That is, these sites show intermediate methylation levels in the majority of individuals. However, in a diploid individual, 50% methylation is only possible if a single copy of the genome is methylated at that site. This follows from assuming unbiased sampling of both copies and the fact that the count data is strand-specific, that is 50% methylation applies either to the positive or negative DNA strand in an individual. Hence, sites with low JSD and intermediate methylation tend to segregate within rather than among individuals. From the count data alone, however, we cannot conclude whether methylation always applies to the maternal or paternal copy, meaning that diversity among individuals is still possible in the narrow sense that mono-allelic methylation is random. In summary, the high proportion of mono-allelic methylated sites among MMCs indicates a substantial heterozygosity of the methylome with respect to the non-CG context. In contrast, the proportion of sites that segregate among individuals (MSCs), being very small in general, is especially small in the non-CG context for rosettes and roots (see Table S3). The proportion of non-CG MSCs is similar to that of CG-MSCs in other sources like embryo, endosperm, and flower bud methylomes but the corresponding sample sizes are small (see Fig. 1c) and we cannot exclude sampling bias or noise effects.

#### Heterochromatin is enriched in CG-HMCs and CHG-MMCs

The chromosome tracks for the proportions of C types in rosettes (Fig. 2d, see Fig. S2 for all sources) show increasing proportions of methylated sites in regions that are rich in repetitive elements and usually heterochromatic: the pericentromeric regions and, for example, the knob in the left arm of chromosome 4. The chromosome arms dominated by protein-coding genes show enrichment for LMCs. This pattern of methylation is in accordance with well-established findings for *Arabidopsis* [3, 4]. However, we can observe that pericentromeric chromatin is dominated by CG-HMCs and CHG-MMCs and that differences between the methylome sources are mainly due to shifts between HMCs and MMCs. In the CG context, HMCs are dominant in all sources but the endosperm. In the CHG context, HMCs are virtually absent in roots and rosettes, slightly increased in embryo, endosperm, and sperm cells, and on a par with MMCs in the vegetative nucleus and flower bud. In the CHH context, HMCs are virtually absent in all sources. MMCs form the smallest fraction of (partially) methylated sites in all sources and are virtually absent in sperm cells. The lack of MMCs in the haploid sperm cells concurs with our interpretation of MMCs given above since heterozygosity is not possible in haploid cells. MSCs also form only a tiny fraction of Cs in the heterochromatic regions with no substantial difference between the sources, although MSCs appear to be slightly increased in endosperm. In contrast to the regions dominated by heterochromatin, there are no regions in the chromosome arms that have a particularly high proportion of MSCs. In conclusion, the enrichment of metastable sites in heterochromatin suggests that this is due to the neutrality of single cytosine polymorphisms in transcriptionally silenced regions.

#### The source determines JSD in the non-CG context

Although genome-wide there is a positive correlation between MET and JSD, we have seen that these quantities can become uncoupled in certain regions of the phase plane and, thus, reflect different properties of the methylome. Hence, we assumed that comparing all combinations of source and context by MET or JSD may lead to different results. By correlating the mean at non-overlapping, 10 kb genomic intervals, we have performed a hierarchical clustering for JSD (Fig. 2e) and for MET (Fig. S3). First, for both genomic signals there is a clear separation into CG and non-CG context. In the non-CG context, however, the MET signals cluster differently than the JSD signals; while context still separates the signals before the source in the case of MET (with the exception of embryo and endosperm, which cluster by source first and then by context, respectively), JSD separates by source first and then by context. Here the exceptions are root, vegetative nucleus, and sperm cell: all three cluster together in the CHG context but in the CHH context, only vegetative nucleus and sperm cluster together, while roots are more similar to rosettes. We think that the general pattern observed here reflects the interplay between maintenance and *de novo* methylation.

There is a robust mechanism in place to maintain CG methylation across cell divisions [25], such that the strong similarity across tissues and cell types is plausible. On the other hand, it has been observed that changes in non-CG methylation accompany cell differentiation, which suggests the effect of *de novo* methylation [26]. In contrast to the strong separation into CG and non-CG context, the merely weak separation into CHG and CHH context along the MET coordinate is overridden along the JSD coordinate through a clear separation into different sources. This may reflect the importance of cell or tissue differentiation, and thus *de novo* methylation, for the non-CG context.

### Chromatin state and methylation diversity

DNA methylation depends not only on DNA sequence features but also on the local state of chromatin. Prominent among the determinants of chromatin state are the location of nucleosomes and the combinations and modifications of their constituent histone proteins. These chromatin marks can interfere with higher-order organization that generates proximity in three dimensions between regions that are distant in the genome. In concert with other regulatory proteins, all of these chromatin marks effect a compaction or relaxation in certain segments of the genome that supports or counteracts the silencing of genes.

In this section, we want to investigate whether the chromatin state correlates with methylation diversity; more specifically, we ask how much diversity in DNA methylation certain regions can tolerate without their chromatin state being affected.

#### Inaccessible chromatin accumulates methylation polymorphisms

The distinction between eu- and heterochromatin is not sufficient to characterize the diversity of chromatin states present in the *Arabidopsis* genome. In order to compare methylation diversity with annotated chromatin features, we used a comprehensive classification of regions into chromatin states based on a multitude of chromatin marks. Sequeira-Mendes *et al.* [29] have identified nine different chromatin states based on DNA methylation, nucleosome occupancy, presence of different histone variants and modifications, and transcriptional activity.

Figure 3a summarizes the profiles of the arithmetic mean of MET and JSD across regions (and 2 kb flanks up- and downstream) distinguished by chromatin state for the rosette. The findings for rosette leaves illustrate by and large a general pattern but some deviations are observed for the heterochromatic states in certain reproductive tissues, e.g. the vegetative cell of pollen and the endosperm of seeds (Fig. S4). Here, we focus on the rosette profiles in the CG context since only the heterochromatic states 8 and 9 show a modest increase in MET and JSD in the non-CG context.

**Figure 3.**
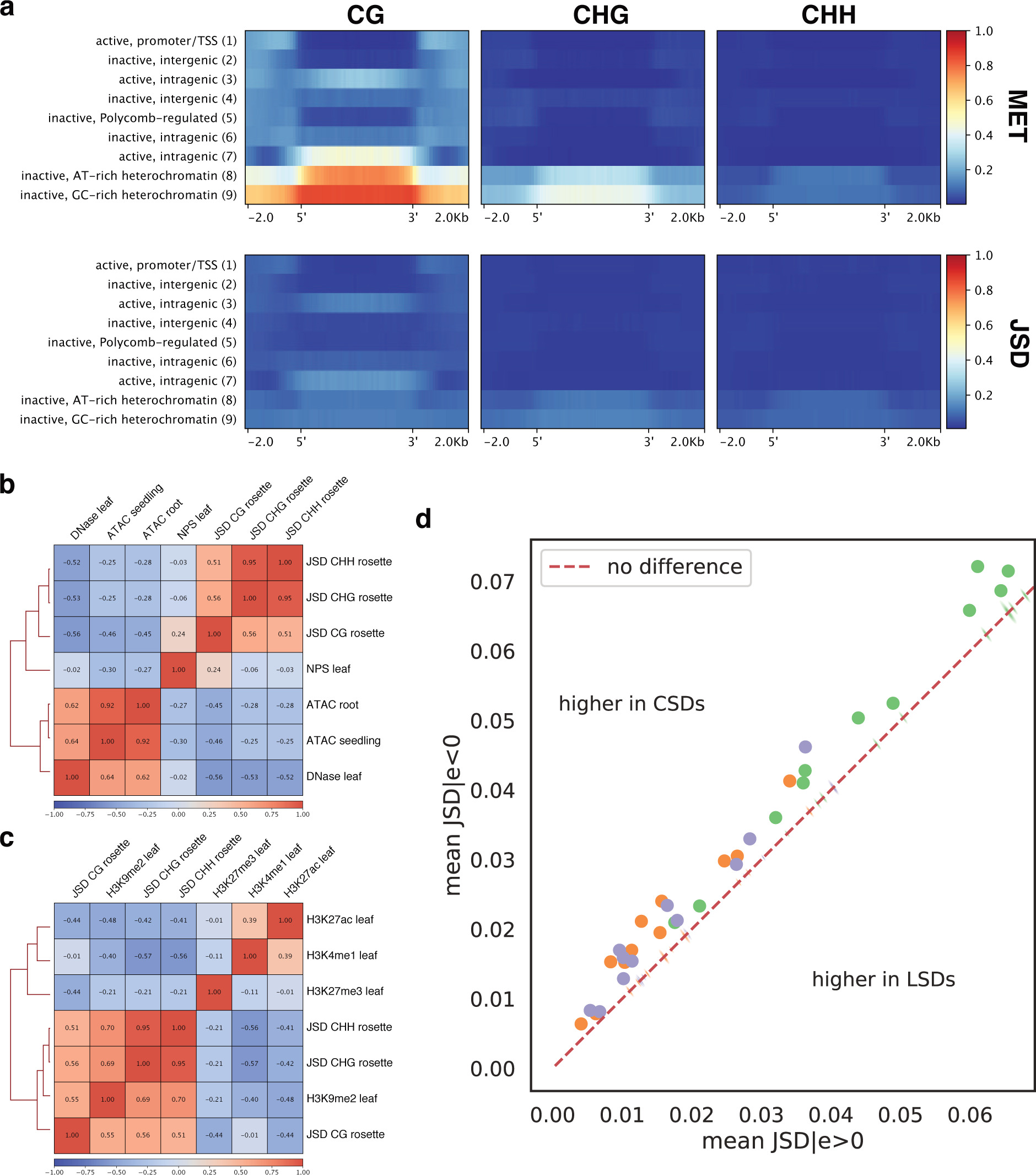
The influence of chromatin state on methylation diversity. **a** MET (top) and JSD (bottom) profiles over regions defined by different chromatin states for rosettes. The profile comprises 2 kb flanks and color-codes the arithmetic mean in non-overlapping, 50 bp bins. **b** Hierarchical clustering of JSD (rosette) with chromatin accessibility signals using Spearman’s correlation coefficient over non-overlapping, 50 kb intervals. Wherever applicable, replicate signals have been averaged. **c** Hierarchical clustering of JSD (rosette) with histone marks. The H3K9me2 signal is log_2_(H3K9me2*/*H3) as in [27]. **d** Mean JSD in regions characterized by positive (*x*-axis) and negative (*y*-axis) HiC-eigenvalues (*e*). The dots indicate the observed pair of values for different C contexts (color-coded) and all sources (not annotated). The kernel density estimates (small ellipses) represent the corresponding random distributions for reshuffled genomic bins. The dashed bisecting line divides the plane into regions of higher JSD in compacted (CSDs) and loose (LSDs) structural domains as defined in [28].

As expected, MET levels are highest in the heterochromatic states 8 and 9, followed by the transcriptionally active, intragenic states 3 and 7 and the inactive states 4 (intergenic) and 6 (intragenic). The increased MET level in these states translates into increased JSD levels except for the intergenic state 4. The JSD levels in the active states 3 and 7 tend to be similar to those in the heterochromatic states while the inactive state 6 shows lower JSD. If we focus not only on the levels in a specific region but on the whole profile, including boundaries and flanking regions, we see further differences among the intragenic as well as the heterochromatic states with elevated JSD. In state 3, MET shows a small dip close to the boundaries, increasing again within the flanking regions, and JSD is slightly higher than in the flanks. In state 6, however, MET actually drops and JSD barely differs compared to the flanks. In contrast to both state 3 and 6, a clear upsurge in MET and JSD is observed for state 7. This state shows a MET profile similar to those of the heterochromatic states 8 and 9. However, for state 8 (AT-rich) JSD drops around its boundaries, which is not observed in state 9 (GC-rich).

According to Sequeira-Mendes *et al.* [29], the heterochromatic states have a high propensity for DNAse1-inaccessible sites. Their increased JSD suggests that inaccessible regions tend to harbor more methylation polymorphisms than accessible regions. This is confirmed by a negative correlation of JSD with DNAse1- [30] and ATAC-seq [31] signal levels (Fig. 3b, see Fig. S5 for MET). In accordance, nucleosome positioning (NPS) shows a positive correlation with CG-JSD; however, the correlation is rather weak and even absent in the non-CG context. Accessibility also seems to affect the diversity of intragenic state 7. This state, which is usually located in gene bodies, shows the sharpest increase in JSD among the intragenic states and tends to have more inaccessible sites than other intragenic states. The increase of JSD in the states close to the coding sequence (3) and the transcription termination site (6) seem to be unrelated to accessibility; in fact, state 3 is the most accessible state. Hence, inaccessibility alone cannot explain the intragenic increases in JSD and we have to consider how chromatin state is determined at the underlying level of histones.

#### Histone signatures characteristic for gene bodies and with depletion of activating marks increase JSD

We have seen that state 3, 6, and 7 are the only intragenic chromatin states that show a noteworthy increase in MET and JSD in the CG context. It is therefore reasonable to investigate whether the differences between these states mirror distinct combinations of histone marks. With respect to levels of JSD in the different intragenic states (*S*) we observe the ordering (*S*7 *> S*3 *> S*6). What are the differences between the intragenic states in terms of histone marks that could explain this order?

Let us first highlight common features. All three states are enriched in H3K4me1 and are depleted in H3K27me3. H3K4me1 is a chromatin mark typical for gene bodies. H3K27me3 is a *Polycomb* mark typical for intergenic, repressive chromatin enriched in chromatin states 2, 4, and 5. At genome-scale (Fig. 3c), H3K4me1 shows a substantial negative correlation with JSD in the non-CG but not in the CG context. H3K27me3 shows the opposite pattern, a negative but rather weak correlation with JSD in the non-CG context but a substantial negative correlation with JSD in the CG context. These findings suggest that histone combinations characteristic for gene bodies are conducive to increased methylation diversity in the CG context. Gene bodies are also characterized by a lack of the histone variant H2A.Z, which accumulates close to the TSS [32]. Thus, we would expect increased methylation diversity with increased gene body likeness, that means lower H2A.Z levels. Indeed, for H2A.Z the order is *S*6 *> S*3 *> S*7, the level in state 3 representing the genomic average [29]. This is the exact reverse of the order with respect to JSD, confirming that histone signatures typical for gene bodies correlate positively with CG-JSD.

With respect to enrichment in activating histone marks, we observe the order *S*3 *> S*7 *> S*6 [29]. The lowest transcribed state 6 lacks the common activating marks H3K36me3 and H3K4me(2/3). The highly expressed but partially inaccessible state 7 does have high levels of the activating mark H3K36me3 but harbors only average H3K4me2 and even reduced H3K4me3 levels. Finally, the highly expressed, highly accessible state 3 contains all three activating marks. An activating histone modification that was not included in the chromatin state classification is H3K27ac, the antagonist to the *Polycomb* mark H3K27me3. It is mainly found in gene bodies and correlates positively with gene expression [33]. At genome scale, H3K27ac is negatively correlated with JSD in all three contexts (Fig. 3c).

In summary, our findings suggest that intragenic regions display increased methylation diversity if they have a histone signature typical for gene bodies and are depleted in activating histone marks.

#### Compacted structural domains show increased methylation diversity

The state of chromatin can also be defined based on the three-dimensional architecture of the chromosomes. In particular, Hi-C studies in *Arabidopsis* have revealed that the genome can be segmented into loose and compacted structural domains (LSDs and CSDs, respectively) [28]. LSDs show a high frequency of interactions with distal domains, while CSDs have a high frequency of local interactions. There is some evidence that CSDs represent a more repressive chromatin state.

We used the quantitative representation of this domain structure, the eigenvalues (*e*) associated with the principal component analysis of the Hi-C correlation matrix, to compare it against MET and JSD. While the magnitude of the eigenvalue does not have a biological meaning, its sign has been shown to indicate LSDs (*e >* 0) and CSDs (*e <* 0), respectively [28]. The sign of the eigenvalue has been identified previously in non-overlapping segments of 50 kb length. We have used this observed segmentation to quantify MET and JSD in LSDs and CSDs, respectively. Then we compared the average difference between these domains in the observed to that in randomly reshuffled segmentations (1,000 random sign permutations) and computed an empirical estimate of the p-value. In this case, the p-value is defined as the probability to randomly obtain an average difference of the signals between CSDs and LSDs at least as extreme as the observed one, *P* (〈*S*_*e*<0_〉 − 〈*S*_*e*>0_〉 > 0 | *H*_0_), where *S* denotes the genomic signal (MET or JSD, respectively) and *H*_0_ is a random segmentation where the number of bins with positive and negative eigenvalues, respectively, is constrained to be the same as in the observed segmentation.

In line with the results obtained using the nine chromatin states, we found that both MET and JSD are increased in CSDs, corresponding largely to repressive chromosome domains (Fig. 3d and Fig. S5b). Although the difference between CSDs and LSDs is small in magnitude, it is highly unlikely to be expected by chance; the empirical estimates of the p-value are zero (Table S4) and the distributions associated with the 1000 randomly reshuffled segmentations are very sharp and falling on the bisecting line in Figure 3d that indicates no difference between LSDs and CSDs.

We conclude that compacted chromatin domains are prone to increased methylation and tolerate higher methylation diversity than loose chromatin domains.

### Methylation diversity of genomic features

In this section, our focus is on methylation diversity in protein-coding genes and different TE categories. We want to analyze whether genes with high diversity have common functional or positional features, which could explain higher JSD in general and enrichment with metastable Cs (MSCs) in particular.

#### TEs can provoke higher methylation diversity

TEs are an important factor in sequence evolution as well as major targets of DNA methylation for the purpose of silencing [6, 34]. Silencing may not have accuracy at the single-base level, in which case we would expect TEs to display methylation polymorphisms.

Figure 4a shows metagene profiles of MET and JSD over protein-coding genes. We have subdivided them into genes that are within TEs (TEGs) and those that are not (non-TEGs). The non-TEGs are further divided into genes that have no overlap with TEGs and those that have an overlap with TEGs.

**Figure 4.**
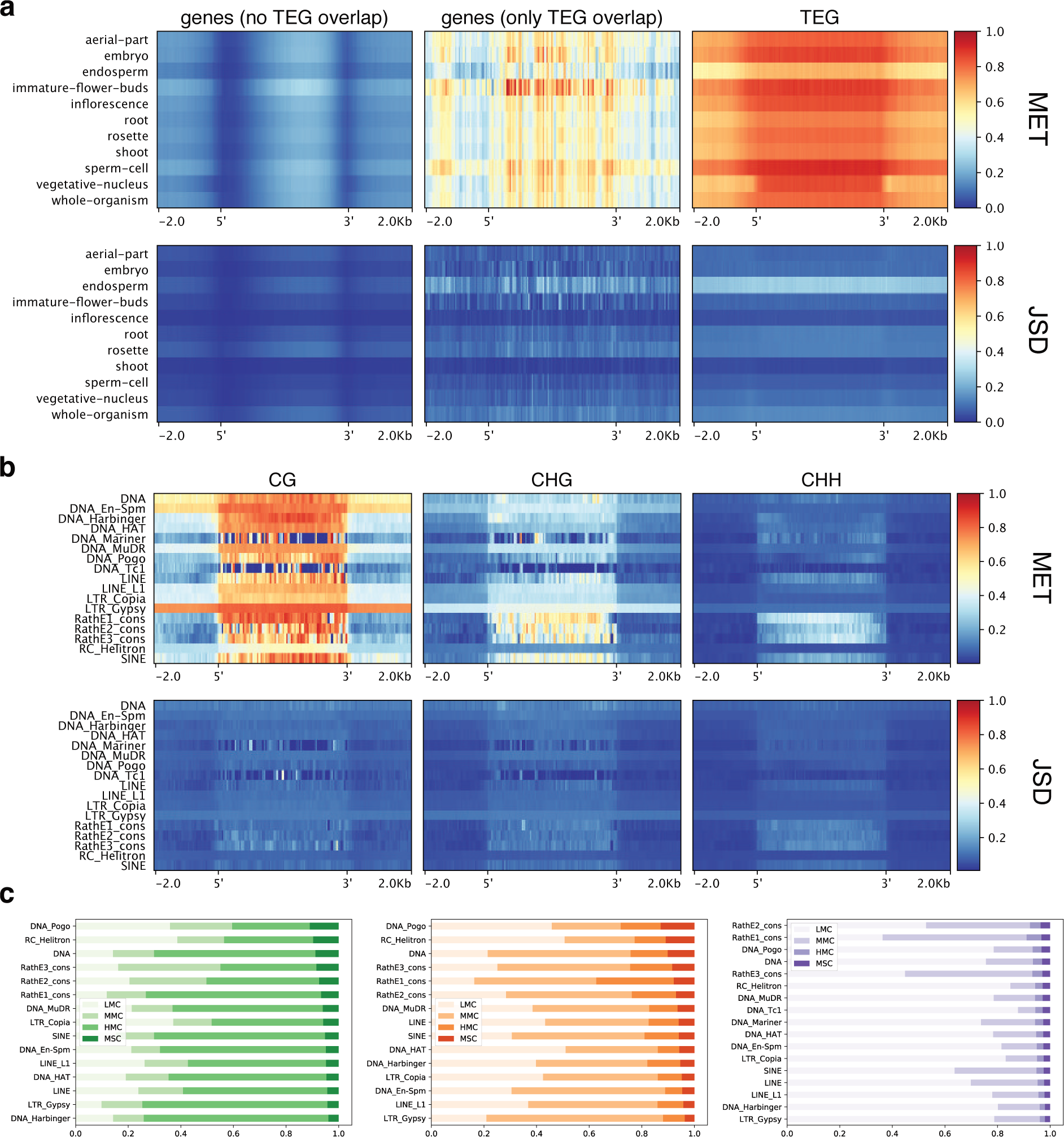
MET and JSD for genes and TEs. All metagene profiles use the arithmetic mean across genomic features with 2kb flanks divided into 50 bp bins. **a** CG-MET and CG-JSD profiles in each source across non-TE genes without TEG overlap, non-TE genes with TEG overlap, and TEGs only. **b** MET and JSD profiles for rosette leaves in each context across TE superfamilies. **c** Proportion of C types for TE superfamilies in each C context sorted by MSC proportion.

Non-TEGs that do not overlap TEGs show the well-known gene body methylation profile for MET in the CG context, with the expected decrease in endosperm. The JSD profile largely follows the MET profile, although the asymmetric bell shape is less pronounced. The non-TEGs with TEG overlap have higher MET and JSD levels than the non-overlapping genes but the TEGs themselves are highly methylated and also show increased JSD levels. Figure 4b shows the MET and JSD profiles of the rosette across all TE super-families in all three contexts. All TE superfamilies display high MET also with respect to their vicinity (DNA Mariner and DNA Tc1 are exceptions that may be due to noise as the coverage of these elements is low). Three superfamilies, DNA, DNA En-Spm, and LTR Gypsy, show increased MET also beyond feature boundaries and, thus, seem to fall into regions that are thoroughly methylated. Most of the TEs have an increased MET in the CHG context as well, but not in the CHH context. The exceptions are non-autonomous retroelements (SINE and RathE(1/2/3) cons super-families) [35], showing increased MET also in the CHH context. The JSD profiles mainly follow the MET profiles. Figure 4c compares the proportions of C types among TE superfamilies in all three contexts, showing that RathE1, RathE2, and RathE3 have rather high proportions of MSCs and MMCs in all contexts, while DNA transposons (DNA, DNA Pogo, and RC Helitron superfamilies) show the highest proportions of MSCs in the CG and CHG context.

In summary, TEs are clearly associated with higher methylation diversity in genes. If TEs elicit DNA methylation without single-base precision, the increased JSD, referring to single-base diversity, in and around TEs makes sense. Note that a consistently methylated region can harbor polymorphisms at the single-base level and show high JSD, even though it was not identified as a differentially methylated region (DMR). This is apparently the case for TEs.

#### Metastable genes are associated with heterochromatic states

TEs are the main drivers of heterochromatin formation. As we have seen, heterochromatic states (chromatin states 8 and 9) have high levels of JSD. Here, we want to investigate whether protein-coding genes that have high proportions of metastable Cs are enriched in heterochromatic regions. Based on that we will take a deeper look in =to the co-localization of these genes with conserved genomic elements that provoke heterochromatin formation.

In the following, we use the term metastable gene (MSG) for genes with a high proportion of MSCs. For each C context, we quantified the proportion of all C types in each gene, followed by sorting the genes according to the proportion of MSCs to select the top 5%. In some sources, this has lead to very low numbers of MSGs and we have excluded these from further analyses. To study the genomic distribution of MSGs without bias, we have normalized the MSG count by the total gene count in non-overlapping, genomic intervals. That is, a high signal in a genomic interval reflects a high proportion of MSGs.

Figure 5 shows the genomic distribution of this fraction of MSGs by context and source over 500 kb intervals. Most of the MSGs are enriched in pericentromeric and telomeric heterochromatin where we usually find a lot of TEs in *Arabidopsis*. However, there are also enrichments outside these regions in the chromosome arms. To see if the identified MSGs are unique, we have analyzed the overlap of MSGs by source and context. The center of Figure 5 shows UpSet plots [36] for overlaps between sources for each context. Interestingly, the MSGs identified with respect to the CG context are mostly unique for each source. We see the opposite in the CHG context, where MSGs are, as a rule, shared among sources; a mixed picture emerges in the CHH context, where we find both unique and shared MSGs. For each source, the overlap among MSGs in different C contexts (Fig. S6) shows that the majority of MSGs are unique for each context.

**Figure 5.**
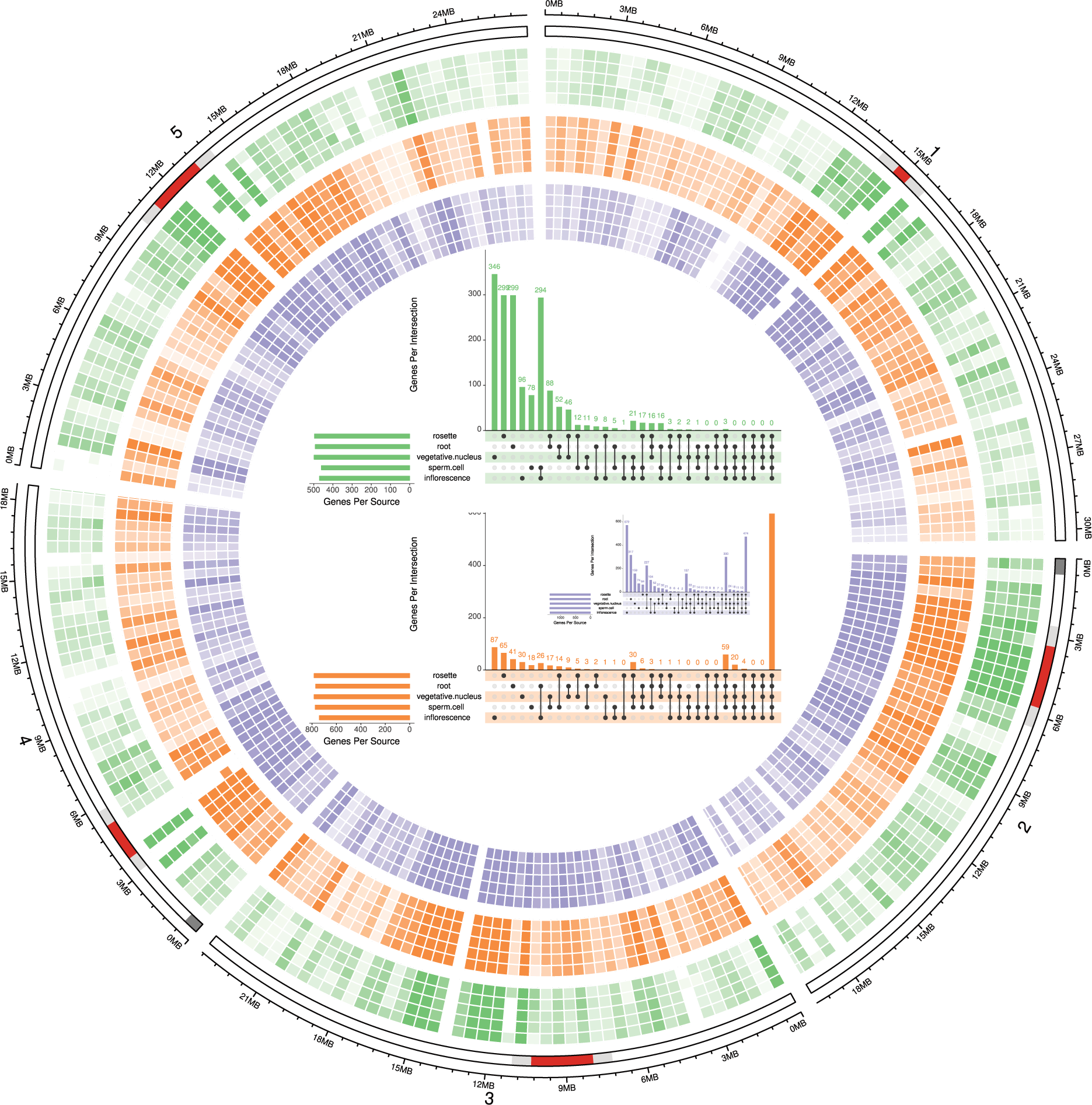
Genome-wide fraction of metastable genes. The axes from outside to inside are chromosome coordinates and idiograms highlighting centromeres (red), pericentromeres (light grey), and telomeric heterochromatin (dark grey). The heatmaps show the fraction of genes that fall into the top 5% percentile of all protein-coding genes (excluding TE genes) with respect to the proportion of MSCs in the open reading frame. The colors encode C context following the colormap used throughout the paper and the opacity encodes the fraction of metastable genes (MSGs). For each context the order of sources from outside to inside is: rosette, root, vegetative nucleus, sperm cell, and inflorescence. **Center**: UpSet plots for MSGs by source in each context. The filled cells below the x axis indicate the sources that are part of the intersection and the y axis indicates the number of MSGs in the respective intersection. The total number of MSGs in each source is given by the bars on the left, respectively.

#### CMT2- and RdDM-targeted TEs can provoke increased methylation diversity in genes

The genomic overview plot in Figure 5 hints at silenced heterochromatin as a determinant of methylation diversity in genes. Silencing often targets TEs to prevent their transposition and mutagenic effects. TEs can be classified into families and superfamilies [35] but also into elements that are targeted by different pathways of the DNA methylation machinery. Based on the analysis of mutants, two groups of TEs have been identified that show differential methylation if either CHROMOMETHYLASE2 (CMT2) or the RNA-dependent DNA methylation (RdDM) pathway is affected [37, 16, 34]; referred to CMT2- and RdDM-targeted TEs hereafter. In this section, we want to explore whether MSGs preferentially co-localize with certain chromatin states and TE categories in order to highlight features that may trigger segregation at the level of DNA methylation.

We looked at the correlation of TE superfamilies, CMT2- and RdDM-targeted TEs, and chromatin states with MSGs to quantify the strength of coenrichment of these features. Figure 6a shows the hierarchical clustering of these elements based on enrichment in non-overlapping, 50 kb intervals. The enrichments are normalized to the count of all protein-coding genes for MSGs, the count of all TEs for the different categories of TEs, and to the length of the interval (here 50 kb) for the coverage with chromatin states, respectively. The distance matrix used for clustering contains the average of Spearman’s correlation coefficient between the enrichments in the intervals. In addition (Fig. 6b), we quantified the spatial correlation of CMT2- and RdDM-targeted TEs with MSGs and chromatin states, respectively, using a relative distance measure [38, 39]. Finally, we performed randomized permutation tests to gauge the evidence in favor of an exceptionally high fraction MSGs close to CMT2- and RdDM-targeted TEs (Fig. 6d).

**Figure 6.**
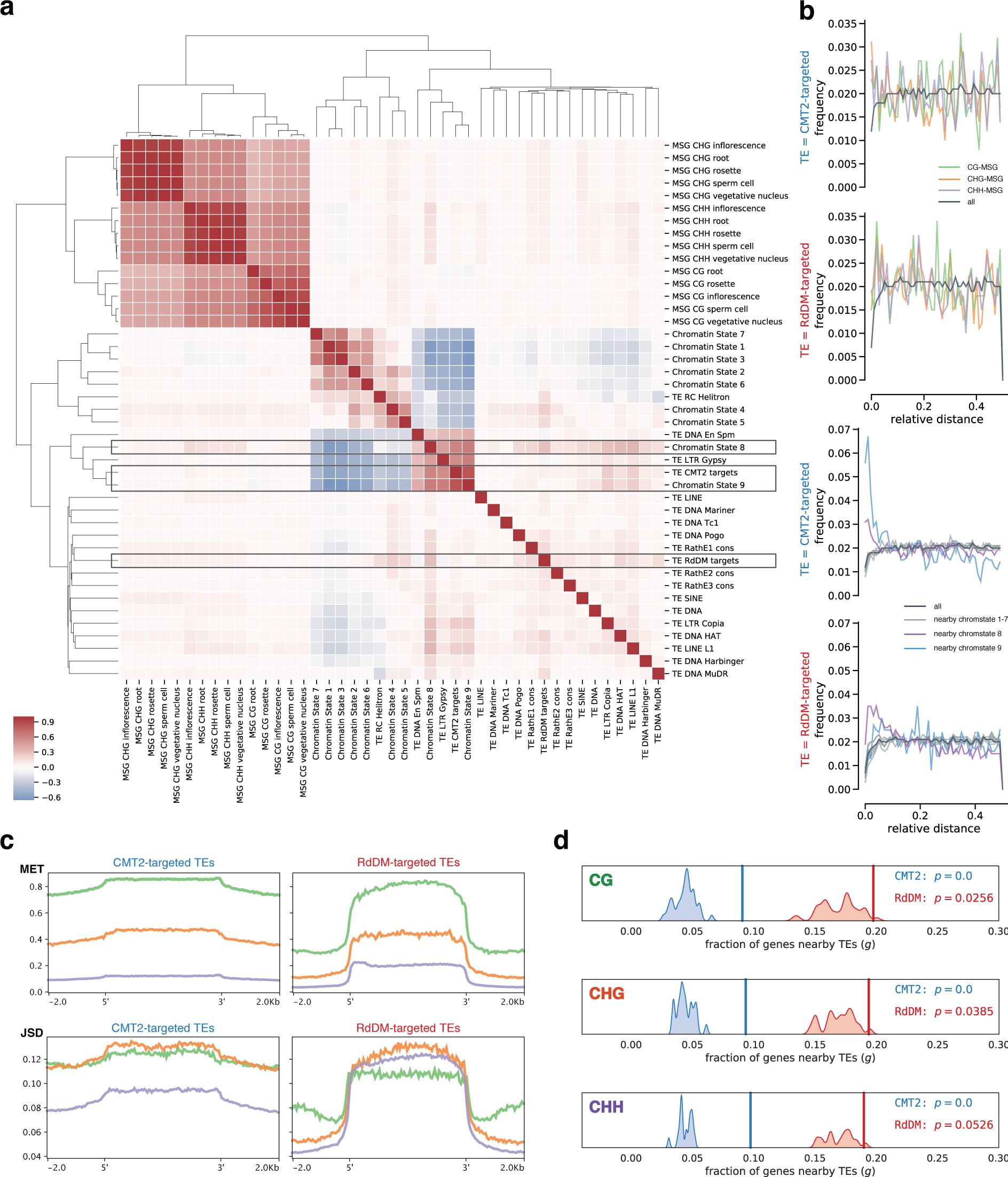
Spatial association of MSGs, TE categories and chromatin states. **a** Hierarchical clustering for normalized enrichment scores of features based on Spearman’s correlation coefficient in non-overlapping, 50 kb intervals. **b** Spatial correlation measured by relative distance between, on the one hand, CMT2- and RdDM-targeted TEs and, on the other hand, genes that are metastable (left) and nearby certain chromatin states (right). **c** MET and JSD profiles for rosette leaves in each context across CMT2- and RdDM-targeted TEs. **d** Probability of metastable genes nearby CMT2- and RdDM-targeted TEs. The *x*-axis gives the gene count normalized to the total number of MSGs (top 5 %, here for rosettes). The vertical lines show the observed fraction of MSGs nearby the targeted TE category. The kernel density estimates show the distribution of the fraction of genes nearby TEs for 10,000 randomly selected gene sets, each of a size equal to the total number of MSGs. The p-value that is estimated by the randomization test is defined by *p* = *P* (*g ≥ g*_observed_ *| H*_0_), where the null hypothesis *H*_0_ is random selection of protein-coding genes that are not TEGs.

According to the hierarchical clustering by enrichment scores (Fig. 6a), there is a clear separation of regions that harbor MSGs from regions that harbor chromatin states and TEs. MSGs cluster by CG and non-CG context as usual. The cluster without the MSGs roughly splits into chromatin states and TEs with some “impurities” in the chromatin state cluster: In the bigger sub-cluster all euchromatic chromatin states (1-7) correlate positively with the RC Helitron superfamily. These TEs are known to be near genes [34]. The smaller sub-cluster, which correlates negatively with the bigger sub-cluster, shows that heterochromatic states 8 and 9 are enriched together with LTR retrotransposons of the Gypsy superfamily, DNA transposons of the EnSpm/CACTA superfamily, and CMT2-targeted TEs. CMT2 preferentially targets Gypsy elements (see Fig. S7) and EnSpm/CACTA elements are known to accumulate along with LTR retro-transposons at pericentromeres, knobs, and TE islands [34]. Both chromatin states and TE superfamilies, respectively, as well as the CMT2-targeted TEs in general show increased levels of H3K9me2 (Fig. S8), the major silencing mark in plants that correlates positively with JSD at genome scale (Fig. 3c). The remaining TE superfamilies and RdDM-targeted TEs form a cluster with rather weak correlation but RdDM-targeted TEs show some co-enrichment with AT-rich heterochromatin (state 8), which itself is co-enriched with MSGs in the CHH context.

The spatial correlation between CMT2- and RdDM-targeted TEs and different groups of genes (Fig. 6b) suggests a close connection between these TEs, heterochromatin, and methylation diversity. Compared to the background of all genes, MSGs are on average closer to CMT2- and RdDM-targeted TEs. The tendency to co-localize with genes near chromatin states 8 and 9 is even more apparent. Interestingly, CMT2-targeted TEs prefer genes near chromatin state 9, whereas RdDM-targeted TEs prefer genes near chromatin state 8. Apart from sequence composition (AT-rich vs. GC-rich), state 8 also differs from state 9 by increased levels of the *Polycomb* mark H3K27me3 and genomic location [29]: State 8 is located in the chromosome arms, interspersed with euchromatic but inactive regions in state 4 (noncoding, intergenic) and 5 (*Polycomb*-regulated), whereas state 9 is characteristic for pericentromeres and is rather interspersed with state 8 only.

The MET and JSD profiles of CMT2- and RdDM-targeted TEs clearly differ (Fig. 6c). It is not so much the level of these signals within the TE boundaries that differs, with MET being highest in the CG, intermediate in the CHG, and barely above noise in the CHH context for both TE categories. In contrast, both signals spread into the vicinity of CMT2- but not RdDM-targeted TEs, since RdDM’s role seems to be to “reinforce the boundary between TE and non-TE” [34]. The phase planes (Fig. S9) are similar in the CG and CHH context with peaks at high and low MET, respectively. In the CHG context, MET is more evenly distributed but reveals a peak at LMCs in RdDM-targeted TEs that is absent in CMT2-targeted TEs.

It seems that metastable genes tend to be close to CMT2- and RdDM-targeted TEs. We tested for this by comparing the observed fraction of MSGs near TEs to the fraction in random gene sets (10,000 random draws of gene sets of the same size as the set of observed MSGs). As Figure 6d clarifies, the distributions for random gene sets are consistently centered below the observed fractions (vertical lines) which makes it highly unlikely that the association of MSGs with CMT2- and RdDM-targeted TEs is by chance. Note that although the fractions for RdDM-targeted TEs are higher than for CMT2-targeted TEs, the statistical evidence is even stronger for CMT2-targeted TEs.

In summary, these results suggest an important role for silenced, heterochromatic elements and associated TEs in driving intragenic methylation diversity. Notably, MSGs co-localize with CMT2- and RdDM-targeted TEs. Therefore, it is likely that TE insertions provoke imprecise *de novo* methylation through CMT2 and RdDM and thereby introduce single methylation polymorphisms that are stabilized through maintenance methylation and amplified into metastable (i.e. segregating) sites characterized by high JSD at the population level.

### Methylation diversity is unaffected by gene expression

Location is not the only factor that affects gene methylation. In fact, a long-standing discussion in epigenetics concerns the interplay of DNA methylation and gene expression. Here, we are interested in the differences in methylation (and methylation diversity) between differentially expressed genes.

To this end, we have profiled MET and JSD in two different categories of genes—the top 50 and bottom 50 genes by relative expression level. Since the expression of genes differs between tissues and organs, we have looked at these two categories in carpels, mature pollen, roots, and rosettes during the vegetative phase of the life cycle using data from the *Arabidopsis* expression angler [42]. Hence, different groups of genes were compared for each transcriptome source. We compared the metagene profiles across tissues, that is for a source-specific pair of gene sets (e.g. top and bottom expressed genes in rosette), we have looked for differences not only in the signals coming from the same source but also in the other sources that where included in this study.

Effectively, there is no difference in methylation between the top 50 and bottom 50 expressed genes of all analyzed sources with one exception: the mature pollen, where the gene bodies of the downregulated genes show on average higher levels of MET and JSD in the CG context (Fig. 7a). The increased diversity in these genes is not restricted to the sperm cell and vegetative nucleus methylomes, which make up the pollen methylome, but is also seen in methylomes from vegetative sources like rosette leaves. Interestingly, many of the genes that are repressed in mature pollen are expressed constitutively in the rest of the plant (see Fig. S10 for a snapshot of the 9 least expressed genes). That means, although there is a difference in methylation between the top 50 and bottom 50 expressed genes in mature pollen, the methylation level of the bottom 50 genes themselves does not change in response to the repression during pollen development. One example is given in Figure 7b. The *AT2G01060* gene, encoding a myb-like HTH transcriptional regulator family protein, shows gene body methylation and is upstream of CMT2- and RdDM-targeted TEs that are associated with high methylation in all three sequence contexts. The inset image shows the repression of this gene in mature pollen.

**Figure 7.**
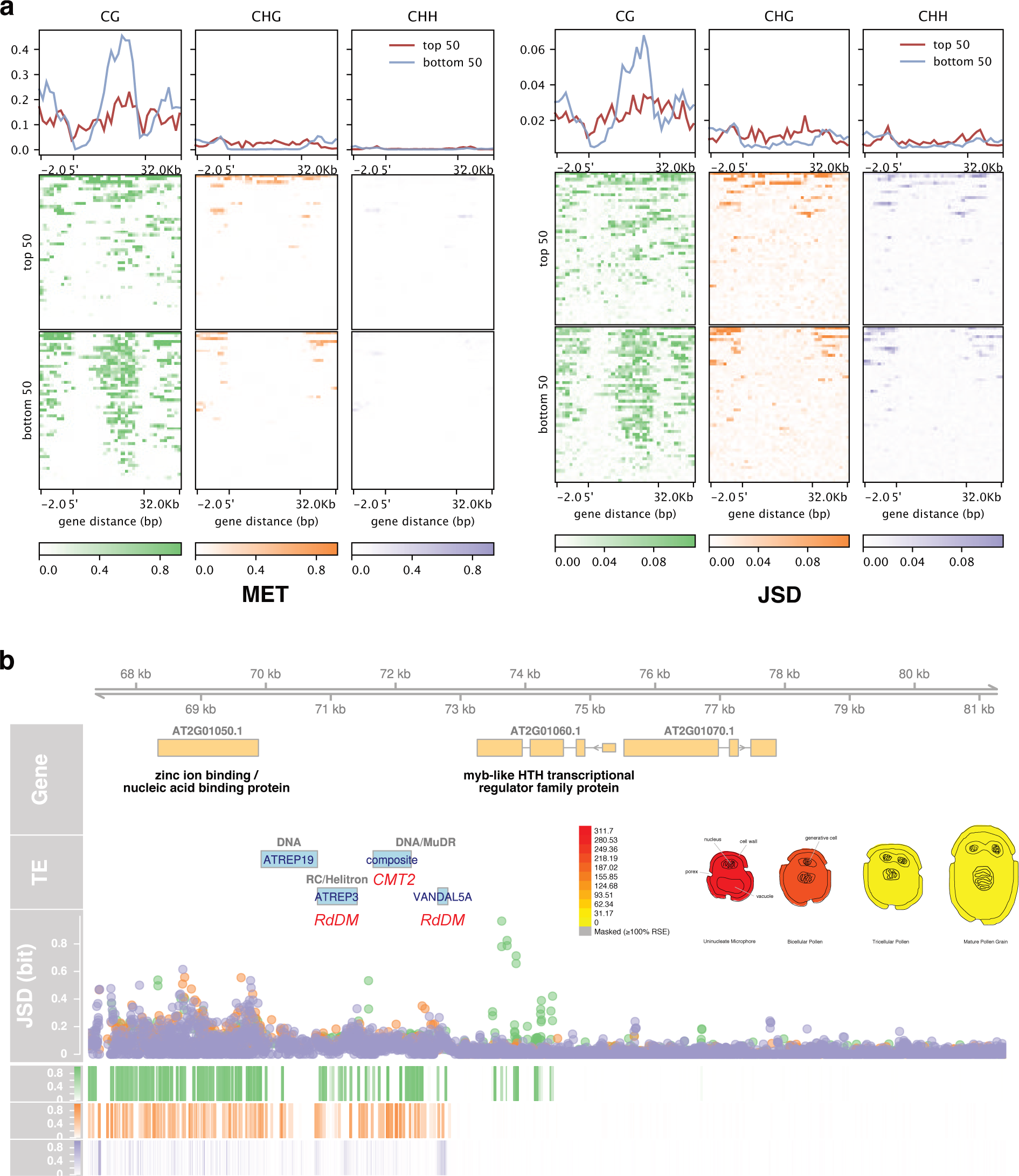
Differential methylation for differentially expressed genes in mature pollen. **a** MET (left) and JSD (right) profiles (arithmetic mean, 2kb flanks, 200 bp bins) from sperm cells for high- (top 50) and low-expressed (bottom 50) genes in mature pollen. The top panels show the metagene profiles over all top and bottom 50 genes, respectively. The bottom panels show profiles of each gene as a heatmap. **b** A genome browser view centered at a gene in the bottom 50 expressed group in mature pollen (myb-like HTH transcriptional regulator family protein; AT2G01060). The TE track highlights the TE family (blue), the superfamily (grey), and the targeting DNA methylation pathway (red) if applicable. The signal tracks are for JSD (scatter plot) and MET (heatmaps) root using context dependent colors as in the rest of the text. The eFP image shows relative gene expression of AT2G01060 and was generated with the “Tissue Specific Microgametogenesis eFP” of ePlant [40] using Affymetrix ATH1 array data [41].

In summary, our analysis of differentially expressed genes across many sources highlights numerous counterexamples to the general claim that variation in gene body methylation is somehow related to expression levels. The increased methylation in the bottom 50 genes expressed in mature pollen is probably a remnant of ancient methylation that is perpetuated through maintenance methylation: first, only CG methylation is different from the top 50 genes, and second, CG methylation does not change during development despite changes in gene expression. Taking also previous negative findings into account [32], a general association between gene expression and gene body methylation seems dubious.

## Discussion

The characterization of the methylome in terms of weighted methylation level (MET) and a diversity measure (here JSD) provides what one may call a *meta-methylome*. Depending on the resolution of the methylomes in the population sample, the meta-methylome can be specific to a species or to different tissue and cell types. In any case, MET has been already identified as the most suitable summary statistic for the methylation level across many sites [43]. It is reassuring that MET turns out to be a by-product of measuring methylation diversity by JSD—together, they define the state of each cytosine in the population. Different approaches based on Shannon entropy already exist to detect differentially methylated sites or regions (reviewed in [44]). However, with the meta-methylome concept, we aim at characterizing the state of a population rather than merely measuring the statistical evidence for differential methylation (although JSD can be used in this way as it generalizes the chisquared test [22]).

This is, to our knowledge, the first application of JSD [21] to DNA methylation. As a symmetrical and smoothed version of the Kullback-Leibler divergence [45], JSD has a solid theoretical foundation and has been successfully applied in many different fields, either as a divergence, as we did, or as a proper distance metric when using the square root of JSD. JSD has some unique properties (discussed in [22, 46, 47, 48]) that set it apart from the multitude of available divergence measures [49]. Interestingly, JSD can also be interpreted as a generalization of Wright’s *F*_ST_, the “most widely used descriptive statistics in population and evolutionary genetics” [1]: the average entropy 〈*H*〉 is the analog to *F*_IS_, the inbreeding coefficient of an individual relative to a subpopulation; the entropy of the mixture, *H* 〈*P*〉, is the analog to *F*_IT_, the inbreeding coefficient of an individual relative to the total population. While *F*-statistics are based on the Pearson correlation coefficient (which has statistical power only in the linear regime), JSD is based on mutual information which can handle any type of statistical dependence [50]. The concept of mutual information has been used to quantify genetic diversity [51, 52]. A serious limitation of the implementation of JSD in its current form, however, is that it only addresses within-population diversity. The extension of JSD to more levels of population subdivision, similar to hierarchical *F*-statistics [1, 53] is outstanding but it would allow us to apportion diversity to different levels. This would increase the practical value of the JSD-based approach to detecting diversity at genome scale.

One way to improve the present approach is to use better estimators for JSD. Certainly, the empirical (“plug-in”) estimator, which replaces probabilities by frequencies, is fast to compute and gives good results for large sample sizes. But better estimators exist for small sample sizes based on the *k*-nearest neighbor algorithm [54], and these may be relevant for estimating JSD at sites with small read depth. Also, the empirical estimator is sensitive to unequal coverage at a site—if one or very few sample units dominate the total read depth at a site, the JSD of the sample will underestimate the JSD of the population. In statistical parlance, the detection of polymorphic sites is prone to false negatives for unequal read depths, an instance of a common problem in statistics when dealing with unequal sample sizes. However, the bias towards false negatives rather than false positives is a tenet for exploratory data analysis. At genome scale, we want to be conservative in our selection of candidate sites rather than following many potential dead ends. Apart from developing and implementing better estimators for JSD, it is also useful to quantify their uncertainty. The uncertainty estimate can be in the form of (orthodox) confidence or (Bayesian) credible intervals. For example, if the sample size is small yet representative, resampling techniques like the bootstrap or the jackknife are appropriate to compute confidence intervals for estimators in the absence of exact formulae [55]. Alas, the determination of uncertainty with randomized algorithms is computationally expensive. A future task is to test if uncertainty quantification is feasible at genome scale.

The diversity analysis of *Arabidopsis* underscores the remarkable overall stability of the methylome across different conditions and supports the conclusion that methylation patterns are to a significant extent determined by genome organization and not by environmental impacts [56, 57]. In regions that show a stable methylation state (i.e. hypo- or hypermethylated regions), JSD can uncover whether precision at the single nucleotide level is critical. If the region shows elevated JSD across the population, as is often the case in heterochromatin, the state of each single cytosine is less important than the state of the region itself. The consideration of different organs, tissues, and cell types highlights some features that may be overlooked if one focuses only on the whole organism or a single tissue: some loci are controlled by gene regulatory processes that unfold during development; in the non-CG context, mainly affected by *de-novo* methylation, patterns of variability are determined by source rather than the difference between the CHG and CHH context. Thus, intra-individual data pooling can indeed obscure inter-individual differences. The same conclusion has been drawn recently in other plants [58, 59] and mammals [60, 26].

This study has shown, that gene body-like chromatin signatures correlate with increased methylation diversity in the CG context. Despite the stability of CG methylation, compared to the non-CG context, a stronger segregation in the CG context is to be expected if faithful maintenance follows noisy *de novo methylation*: The positive feedback loop at the heart of CG methylation [25] blows up any minor variation in the original cell population and stabilizes different states leading to metastable cytosines or epialleles at the population level [61]. However, metastable alleles will only survive if there is no selection pressure to weed out detrimental methylation variants. In gene bodies at least, the methylation level of single cytosine sites, although not necessarily that of the whole body, appears to be selectively neutral. Thus, methylation in gene bodies is not precision work but a crude act to shut down deleterious DNA.

One form of deleterious DNA that needs to be silenced are TEs. In *Arabidopsis*, being an organism that uses DNA methylation for silencing, the RdDM and CMT2 pathways can target TEs that are nearby or overlap genes. As we have seen, these silencing mechanisms are probably triggering increased levels of methylation diversity in general and metastable Cs in particular. Our results support and refine previous indications that TEs influence epigenomic diversity [16, 62, 63]. Certainly, random variation in the DNA sequence is the foundation of evolution. “Selfish” genetic elements, though largely deleterious, can become functional [64, 65] and may even play a role in speciation [66]. However, if they provoke a, necessarily noisy, silencing response, an epigenetic layer of variation emerges that may influence the “selective arena”. As much as genetic diversity is driven by “loud” TEs, epigenetic diversity is driven by silent TEs—they become important drivers of evolution even if they appear to be neutralized.

## Conclusions

We have implemented a fast, scalable method to perform genomic scans of diversity in large populations. This approach based on JSD is non-parametric; hence, it works without parameter tuning and model specification. JSD can be applied to any functional genomics data that maps a discrete probability distribution to a locus. Furthermore, its application is general and can be extended to analyze epigenetic variation between individuals, organs, tissues, or cells, including different cell lineages in heterogeneous tumors [8]. The application of JSD to methylome data in *Arabidopsis* shows that methylation diversity tends to increase the more closed, heterochromatic or silenced chromatin is. Our analysis emphasizes the dominant role of location for methylation and its diversity, in particular the putative impact of nearby TEs that are targeted by CMT2 and the RdDM pathway.

## Methods

### Jensen-Shannon Divergence

JSD is an information-theoretic divergence measure based on Shannon entropy [23]. As any divergence measure, it assigns a real number to a set of probability distributions defined on a common sample space. This number reflects the diversity of the set. Formally, the general Jensen-Shanon divergence (*D*) for a distribution set *P* is defined in terms of Shannon entropy *H* as [47]

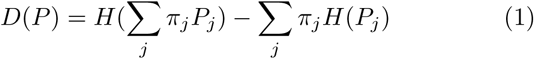

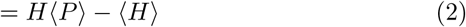

Here, the mixture distribution 〈*P*〉 = Σ_*j*_ *π*_*j*_*P*_*j*_ is the average of the probability distributions *P*_*j*_ with respect to the normalized weights *π*_*j*_, where Σ_*j*_ *π*_*j*_ = 1, and 〈*H*〉 is the corresponding average of the entropy of all *P*_*j*_. The Shannon entropy for a discrete distribution is generally defined as

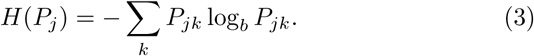

Here, *k* ∈ Ω is an event from the sample space with probability *P*_*jk*_, such that Σ_*k*_*P*_*jk*_ = 1 is fulfilled. Regarding the logarithm, we follow the convention in information theory and use *b* = 2 as the base, such that JSD is measured in *bit*.

From the viewpoint of JSD, diversity is equivalent to the expected loss of information upon mixing. Figure 1a illustrates this geometrically using binary distributions (i.e. with two events in the sample space). The maximum entropy in bit is log_2_(2) = 1 bit in this case. The mixture entropy, *H*〈*P*〉, must lie on the red segment of the entropy graph, the exact location depending on the weights. Likewise, the point representing the corresponding average entropy, 〈*H*〉, of the set must lie on the red triangle. Due to the shape of the entropy graph, the red segment will always be above the red triangle, which means that *H*〈*P*〉 ≥ 〈*H*〉. Essentially, this inequality expresses the expectation that mixing different sources of information tends to increase uncertainty or, equivalently, leads to a loss of information. Due to this inequality, JSD is always bounded by zero and the logarithm of the size of the sample space, 0 *≤ D ≤* log(|Ω|).

### Methylation Diversity

To compute methylation diversity over a reference genome, we have to estimate JSD from a set of methylation tables, that is from read counts for two different events over a collection of methylomes. Let *i*, *j*, and *k* be indices for cytosine position in the genome, methylome in the population sample, and methylation state, respectively. Without loss of generality, we let *k* = 1 indicate the methylated state and *k* = 2 the unmethylated state, such that *n*_*ijk*_ denotes the corresponding read count. Based on these read counts, a straightforward estimate of population JSD is obtained by replacing probabilities with sample frequencies; this leads to the so-called *plug-in* or *empirical* estimator of JSD at each position *i*:

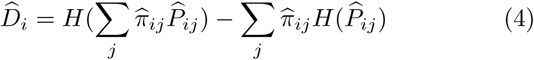

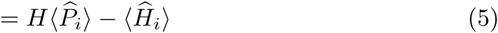

Here, the distributions and weights are replaced by their empirical counterparts

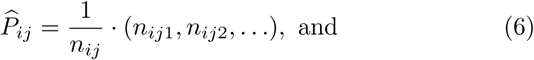

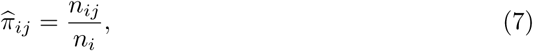

where *n*_*ij*_ = Σ_*k*_*n*_*ijk*_ is the per-methylome coverage at position *i* in methylome *j*, and *n*_*i*_ = Σ_*j*_ *n*_*ij*_ is the total coverage of position *i*. Grosse *et al*. [22] have shown that the plug-in estimator gives the maximum likelihood estimate of JSD if empirical weights are used. Table 1 shows by example how the plug-in estimate of JSD is computed. We have developed an open-source program in Python, tentatively called *Shannon*, with a simple command line interface to efficiently perform JSD scans for a large set of methylomes using the plugin estimator, see Supplementary Section S1.1 for more details.

A by-product of computing JSD at a site *i* is the methylation level 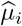, which is the weighted average of the methylation levels 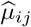 of the sampling units in the population sample:

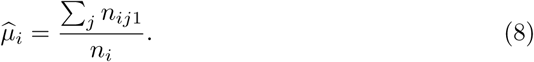

This is the plug-in estimate of the methylation bias (MET) within the population. Unless stated otherwise, we refer to the position-specific estimates 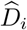 and 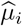 whenever we speak of concrete JSD and MET values. Figure 1b shows that JSD and MET span a “phase plane” that visualizes the spectrum of methylation across the genome and the population. At the population level, each cytosine can be represented by a combination of JSD and MET, hence a point in the phase plane. Since *H* 〈*P*〉 is an upper bound for JSD, all such points must fall within the region below the curve. Depending on the position in the phase plane, one may distinguish different cytosine types (C types) as Figure 1b suggests: LMCs (low-methylated cytosines), MMCs (medium-methylated cytosines), HMCs (highly-methylated cytosines), and MSCs (metastable cytosines that segregate in the population). Any boundary that defines these regions is to some degree arbitrary as it cannot be derived from first principles. However, reasonable methylation level thresholds are 20% and 80% for low and high methylation, respectively. In support of this choice, consider that in Bayesian models for methylation level inference, the beta-binomial distribution is often the natural choice as the likelihood function [67]. The methylation level of the population sample, MET, indicates the peak of the posterior density for methylation; hence MET *>* 0.8 would translate into a high probability of methylation for the given site.

### Data processing

To generate the metadata, custom Python scripts were used to query the European Nucleotide Archive (ENA) and NCBI’s Biosample database for all available *Arabidopsis* BS-seq runs which were subsequently curated semi-automatically to remove inconsistencies. The investigation was restricted to wild-type methylomes of the Col-0 accession and to tissues/organs (hereinafter called source), for which at least three methylomes were available, see the overview in Figure 1c.

Based on the metadata, methylation tables were generated by, first, mapping sequencing runs in fastq format (quality-filtered using TrimGalore!/cutadapt [68] and mapped to the reference genome (TAIR10) using Bismark [69]) and, second, making methylation calls with MethylDackel [70]. The methylation tables were subsequently indexed with tabix [71] to prepare the data for computing JSD with *Shannon*.

For the downstream analysis we used the current genome annotation Araport 11 [72]. The complete pipeline was implemented in Snakemake [73], mainly using the scientific python stack [74, 75, 76, 77], R visualization libraries [78, 79, 36], and tools for genome analysis [38, 80, 81]. Further details are given in the Supplementary Section S1.2. The complete pipeline code is available in a git repository (https://gitlab.com/okartal/meta-methylome).

## Data availability

*Shannon* is available at https://gitlab.com/okartal/shannon.git. The data pipeline is available at https://gitlab.com/okartal/meta-methylome.git. All genomic data, tables, and figures are available in Zenodo (https://doi.org/10.5281/zenodo.3521983).

## Author’s contributions

Conceptualization: ÖK and UG; Methodology: ÖK and UG; Software: ÖK and MWS; Formal Analysis: ÖK and MWS; Investigation: ÖK; Data Curation: ÖK; Writing - Original Draft: ÖK; Writing - Review & Editing: ÖK and UG; Visualization: ÖK; Funding Acquisition: ÖK and UG.

## Acknowledgments

We thank Eriko Sasaki and Magnus Nordborg (Gregor Mendel Institute, Vienna) for providing the list of CMT2- and RdDM-targeted TEs. We thank Stefan Grob (Department of Plant and Microbial Biology, University of Zurich) for providing the coordinates of CSDs and LSDs. This work was supported by the University of Zurich, a Transition Postdoc Fellowship from SystemsX.ch (to ÖK), and grants from the European Research Council and the Swiss National Science Foundation (to UG).

## Supplementary Information

### S1 Methods

#### S1.1 *Shannon* — a command line app for computing JSD

We are currently developing an open-source, command line application for POSIX-like operating systems. We chose Python for its general-purpose features and the availability of mature libraries for numerical computing, especially pandas and NumPy. The source code is available on GitLab (https://gitlab.com/okartal/shannon.git). The application needs two types of data inputs:

- tabix-indexed genome position files (GPFs) for each sampling unit in the population sample and
- a metadata table as a tab- or comma-delimited text file where each row corresponds to a sampling unit

The metadata table must have at least two columns: ID, to specify a unique name for each sample, and URL to specify the location of its GPF. In addition, the metadata can contain columns that further specify each sample (e.g. ecotype, tissue, cell type, etc.). The output is a BED-like file with JSD values for each genomic position. Below is the help message and a usage example for computing JSD at Chr1 where the count data are given in the 5th (mC, methylated base calls) and 6th column (C, unmethylated base calls) of the GPFs, respectively:

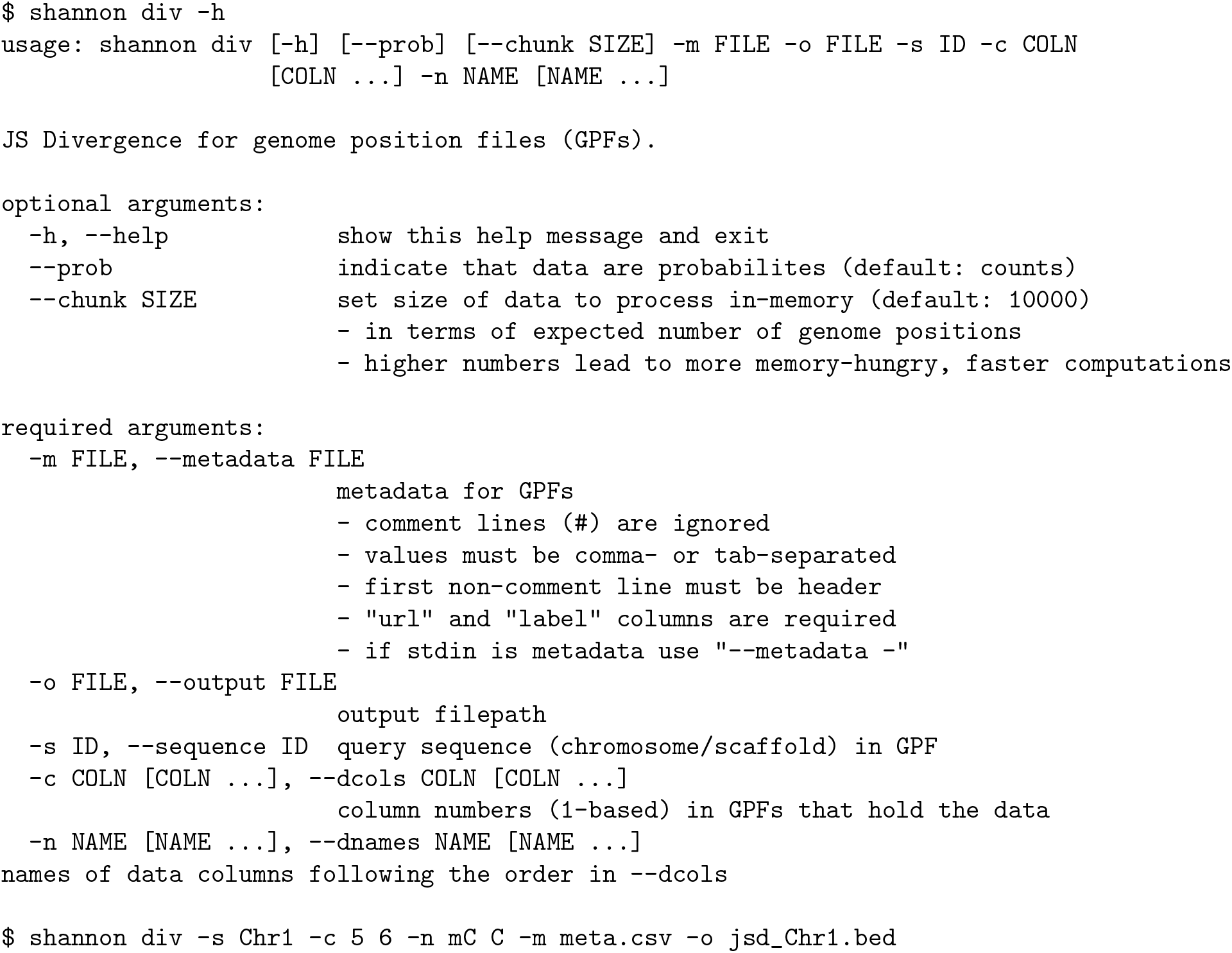

In order to efficiently handle a large number of GPFs, the program chunks the genome into regions and sequentially processes each region by merging data across all GPFs. Reading data relies heavily on fast random access to genomic regions using tabix and efficient collection of the GPF regions using Python generators to prevent I/O-bound slowdowns. Merging data relies on pandas’s ability to concatenate differently indexed data frames and to handle missing values for certain combinations of sampling unit and position. To estimate JSD based on the merged count data, the computations on each chunk are done in-memory using NumPy arrays. As of now, we have only implemented the empirical (“plug-in”) estimator of JSD. That is, we use the empirical distributions to calculate weights, mixtures and entropy terms.

#### S1.2 Software Environment for Analysis

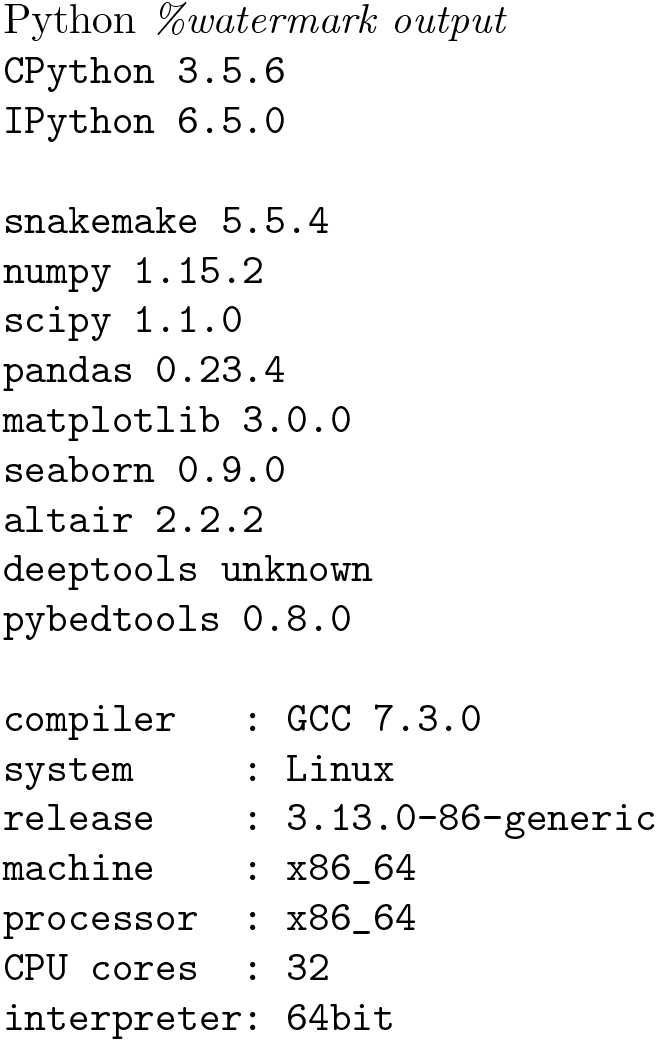

R *sessionInfo()*

- R version 3.5.2 (2018-12-20), x86_64-pc-linux-gnu
- Locale: LC_CTYPE=en_US.UTF-8, LC_NUMERIC=C, LC_TIME=en_US.UTF-8, LC_COLLATE=en_US.UTF-8, LC_MONETARY=en_US.UTF-8, LC_MESSAGES=en_US.UTF-8, LC_PAPER=en_US.UTF-8, LC_NAME=C, LC_ADDRESS=C, LC_TELEPHONE=C, LC_MEASUREMENT=en_US.UTF-8, LC_IDENTIFICATION=C
- Running under: `Ubuntu 14.04.4 LTS`
- Matrix products: default
- BLAS: `/usr/lib/libblas/libblas.so.3.0`
- LAPACK: `/usr/lib/lapack/liblapack.so.3.0`
- Base packages: base, datasets, graphics, grDevices, grid, methods, parallel, stats, stats4, utils
- Other packages: BiocGenerics 0.28.0, circlize 0.4.6, GenomeInfoDb 1.18.2, GenomicRanges 1.34.0, Gviz 1.26.5, IRanges 2.16.0, S4Vectors 0.20.1, UpSetR 1.3.3
- Loaded via a namespace (and not attached): acepack 1.4.1, AnnotationDbi 1.44.0, AnnotationFilter 1.6.0, assertthat 0.2.0, backports 1.1.3, base64enc 0.1-3, Biobase 2.42.0, BiocParallel 1.16.6, biomaRt 2.38.0, Biostrings 2.50.2, biovizBase 1.30.1, bit 1.1-14, bit64 0.9-7, bitops 1.0-6, blob 1.1.1, BSgenome 1.50.0, checkmate 1.9.1, cluster 2.0.7-1, colorspace 1.4-0, compiler 3.5.2, crayon 1.3.4, curl 3.3, data.table 1.12.0, DBI 1.0.0, DelayedArray 0.8.0, dichromat 2.0-0, digest 0.6.18, dplyr 0.8.0.1, ensembldb 2.6.6, foreign 0.8-71, Formula 1.2-3, GenomeInfoDbData 1.2.0, GenomicAlignments 1.18.1, GenomicFeatures 1.34.3, ggplot2 3.1.0, GlobalOptions 0.1.0, glue 1.3.0, gridExtra 2.3, gtable 0.2.0, Hmisc 4.2-0, hms 0.4.2, htmlTable 1.13.1, htmltools 0.3.6, htmlwidgets 1.3, httr 1.4.0, knitr 1.21, lattice 0.20-38, latticeExtra 0.6-28, lazyeval 0.2.1, magrittr 1.5, Matrix 1.2-15, matrixStats 0.54.0, memoise 1.1.0, munsell 0.5.0, nnet 7.3-12, pillar 1.3.1, pkgconfig 2.0.2, plyr 1.8.4, prettyunits 1.0.2, progress 1.2.0, ProtGenerics 1.14.0, purrr 0.3.0, R6 2.4.0, RColorBrewer 1.1-2, Rcpp 1.0.0, RCurl 1.95-4.11, rlang 0.3.1, rpart 4.1-13, Rsamtools 1.34.1, RSQLite 2.1.1, rstudioapi 0.9.0, rtracklayer 1.42.1, scales 1.0.0, shape 1.4.4, splines 3.5.2, stringi 1.3.1, stringr 1.4.0, SummarizedExperiment 1.12.0, survival 2.43-3, tibble 2.0.1, tidyselect 0.2.5, tools 3.5.2, VariantAnnotation 1.28.11, xfun 0.5, XML 3.98-1.17, XVector 0.22.0, zlibbioc 1.28.0

### S2 Tables

**Table S1** Statistics for JSD and MET in all sources: `meta_methylome/new/data/results/stats_Col-0_wt_[SOURCE].csv`

**Table S2** Mann-Whitney *U* test for differences in JSD between C contexts: `meta_methylome/new/data/results/tests-mannwhitneyu_jsd-between-context_Col-0_wt_[SOURCE].csv`

**Table S3.**
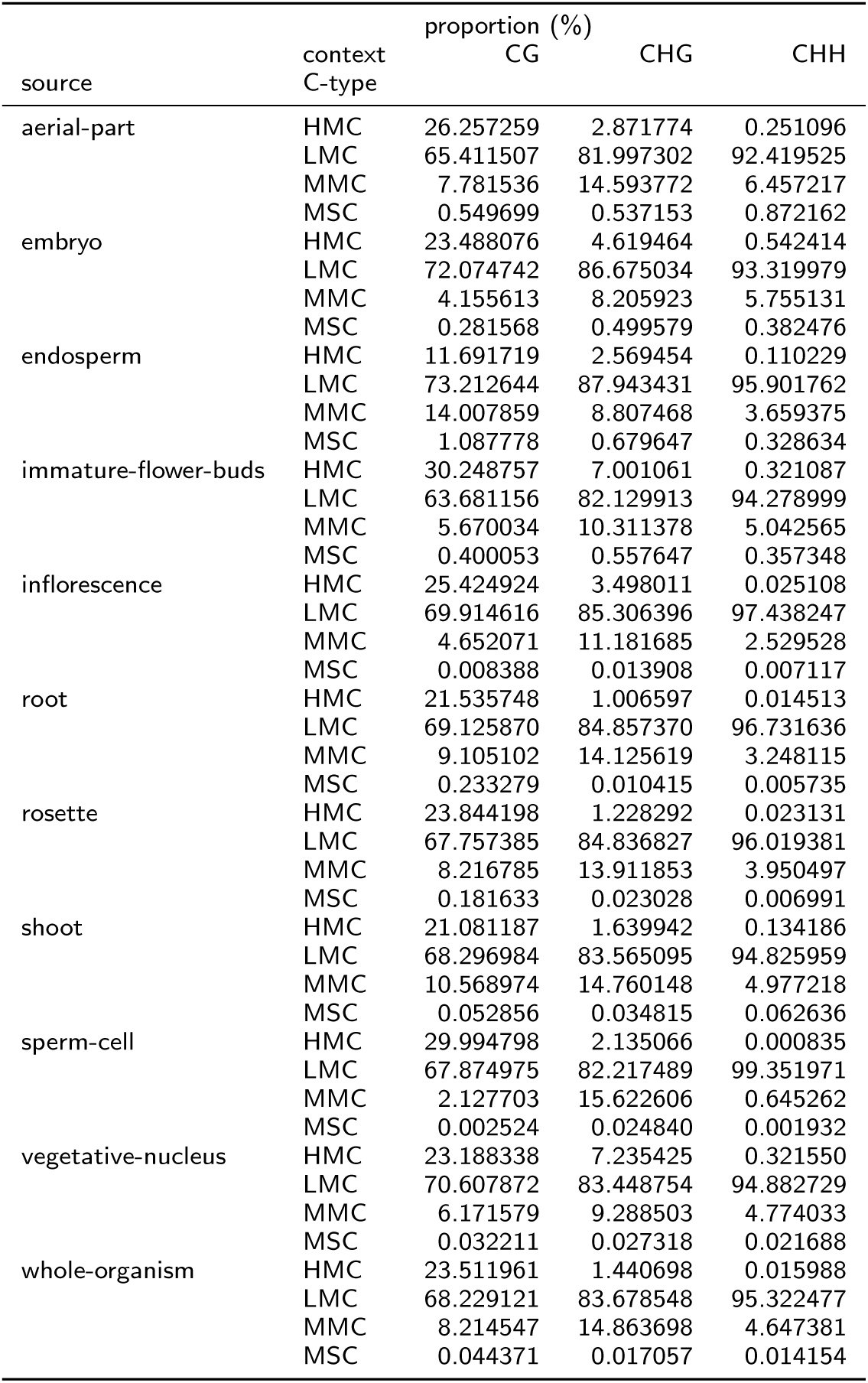
Genome-wide proportions of C types per source and context.

**Table S4** Empirical p-values for getting at least the observed difference between CSDs and LSDs in 1,000 randomly reshuffled segmentations: `meta_methylome/new/data/results/tab-pvalues_HiC-domains_mean.csv`

### S3 Figures

**Figure S1.**
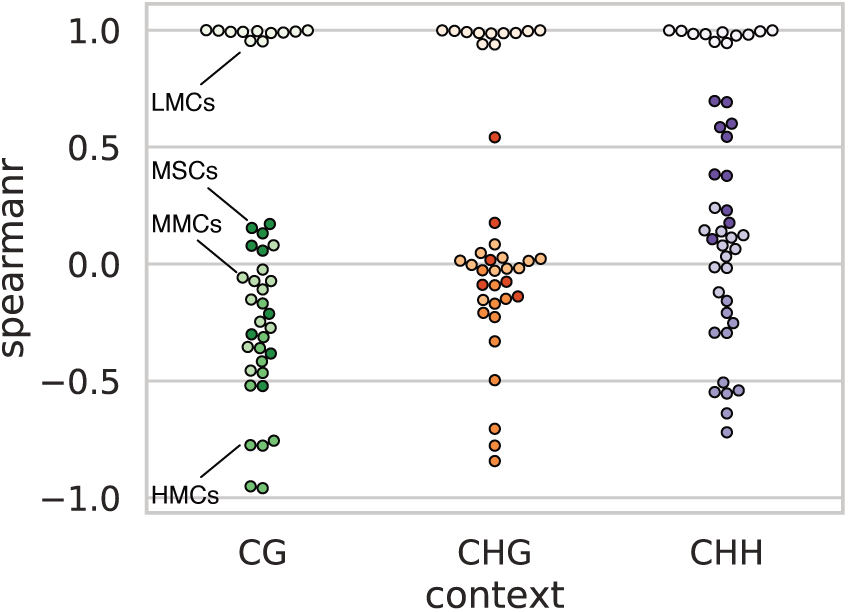
Correlation between MET and JSD. Spearman’s correlation coefficient (y-axis) was measured in each subregion of the phaseplane, defining a C type (encoded by hue), and for each C context (encoded by color and x-axis).

**Figure S2.**
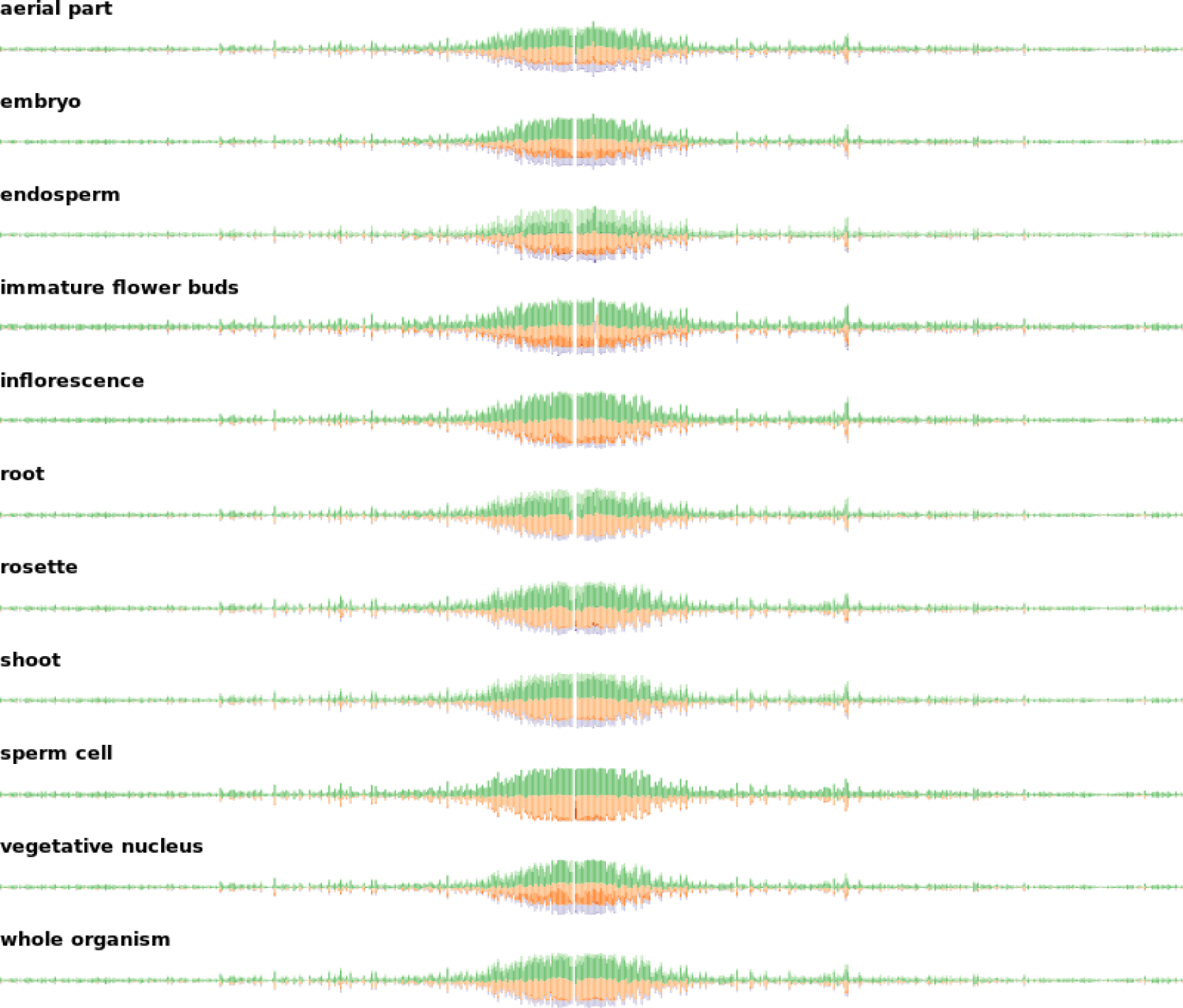
Chromosome 1 tracks for the proportions of C types in all sources. Proportions (stacked on y-axis, respectively) have been calculated at non-overlapping, 50 kb intervals (x-axis). Figures for all chromosomes can be found at `meta_methylome/new/data/results/fig/plot-chromtrack-ctype_[CHROM].png.`

**Figure S3.**
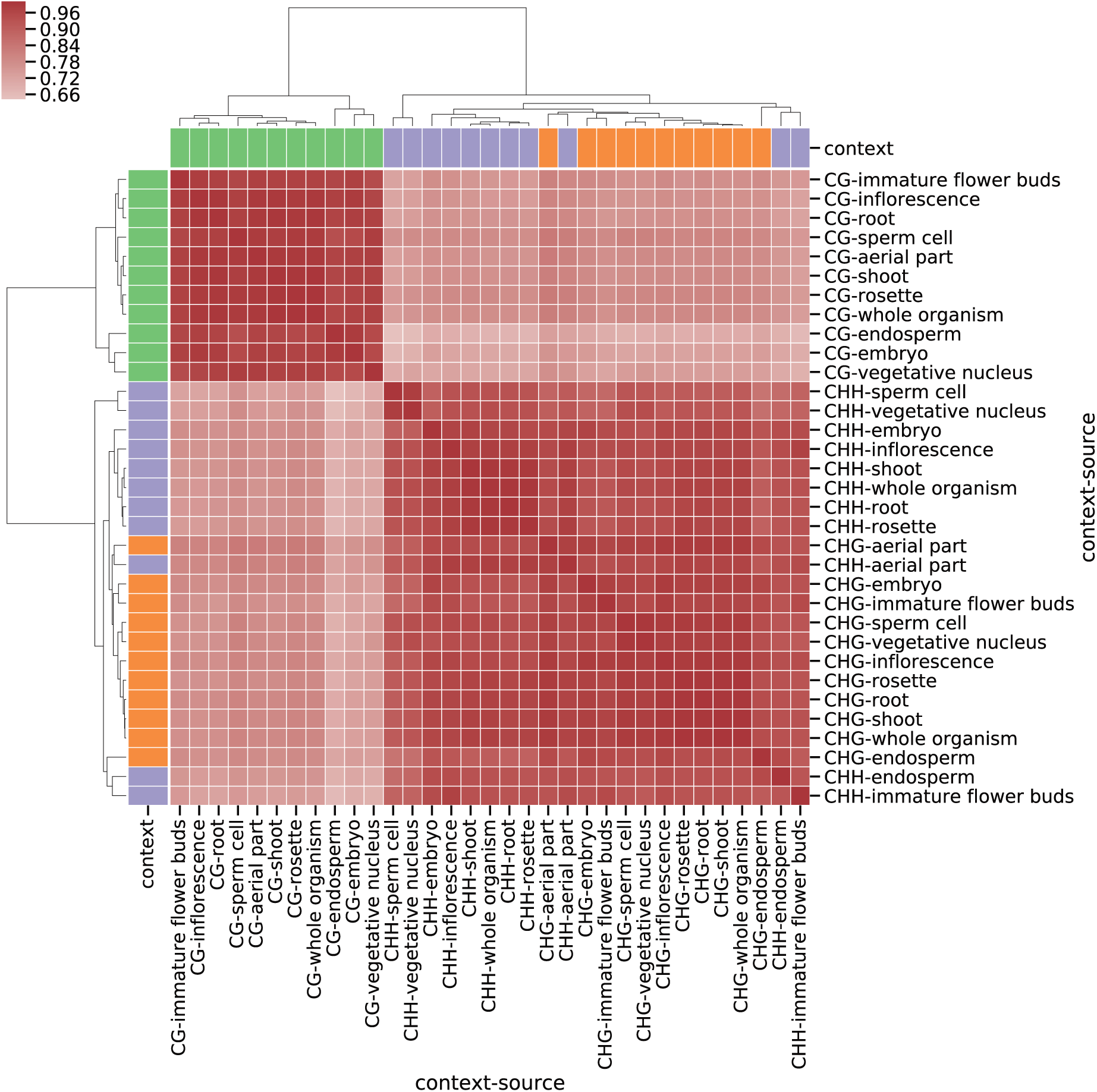
Hierarchical clustering by source and C context using MET. The distance matrix is given by the mean of Spearman’s correlation coefficients at non-overlapping, 50 kb intervals over the whole genome.

**Figure S4.**
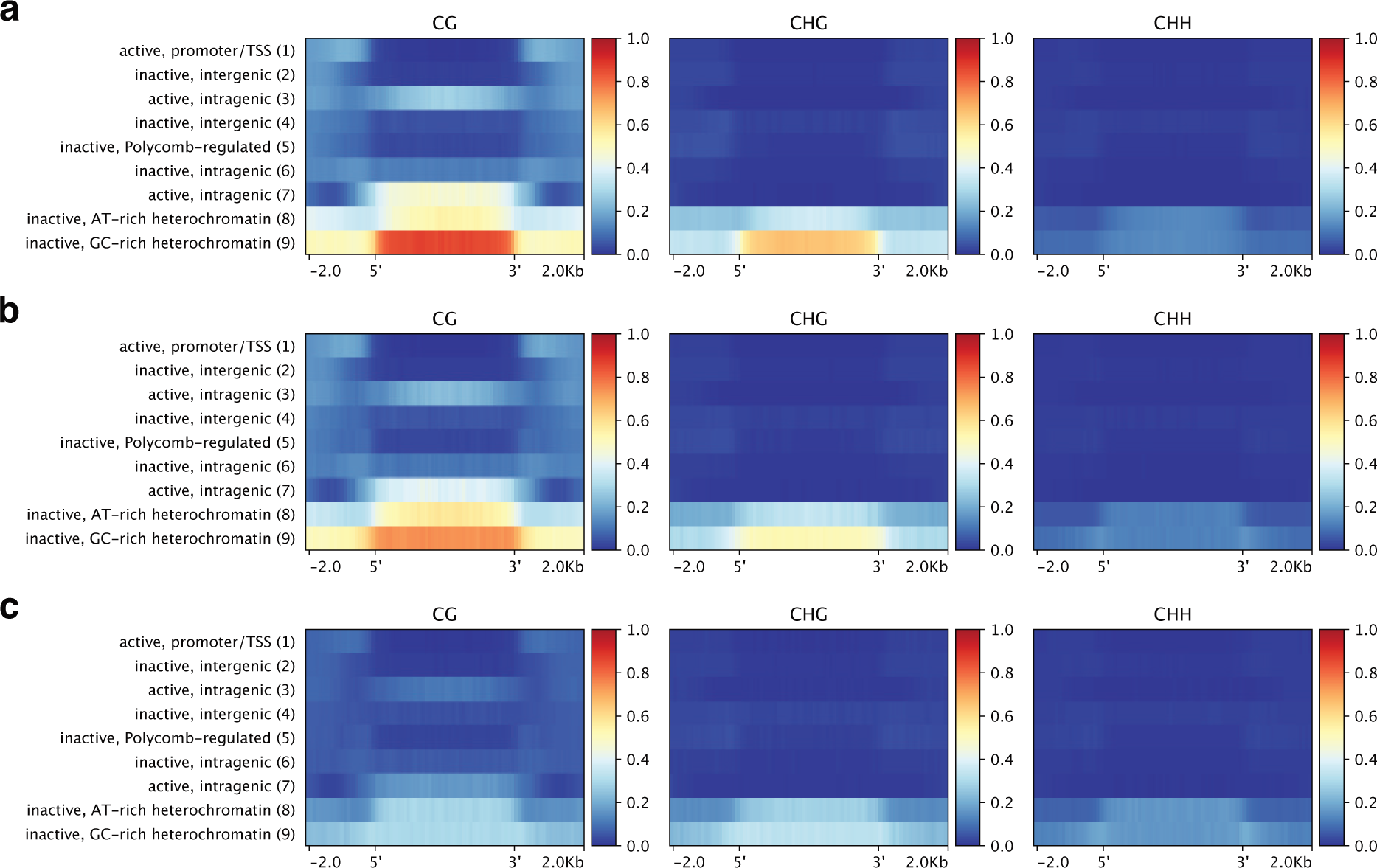
Inuence of chromatin state on methylation diversity. The profile comprises 2 kb flanks and color-codes the arithmetic mean in non-overlapping, 50 bp bins (x-axis). **a** MET in vegetative nucleus. **b** MET in endosperm. **c** JSD in endosperm.

**Figure S5.**
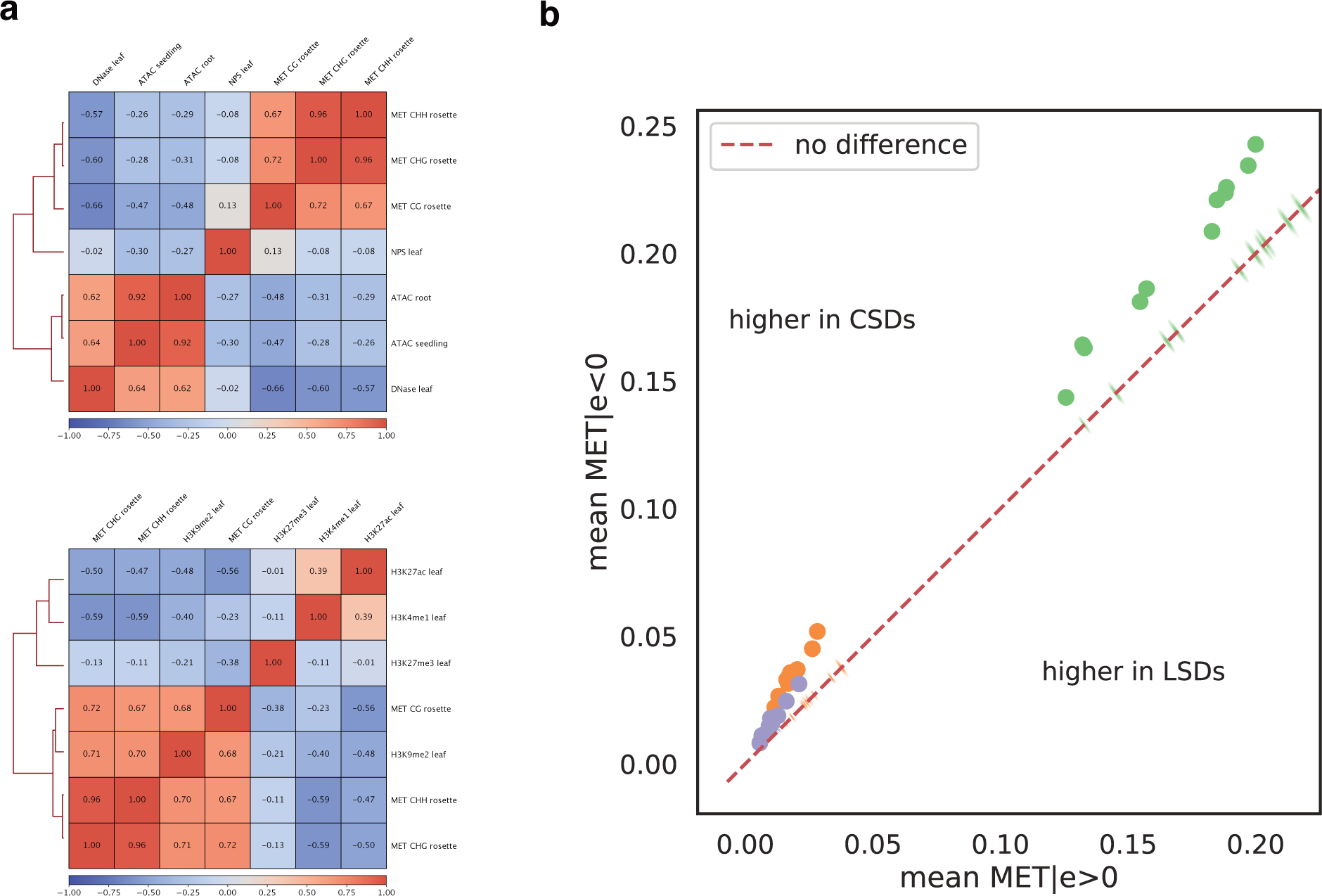
Correlation of chromatin accessibility and histone signals with MET. **a** Hierarchical clustering of MET (rosette) with chromatin accessibility signals (top) and histone marks (bottom) using Spearman’s correlation coefficient over non-overlapping, 50 kb intervals. The number shows the Spearman correlation coefficient averaged over non-overlapping 50 kb intervals. Wherever applicable, replicate signals have been averaged. The H3K9me2 signal is actually log_2_(H3K9me2/H3) as in [27]. **b** Mean MET in regions characterized by positive (*x*-axis, ⋅|*e* > 0) and negative (*y*-axis, |*e* < 0) HiC-eigenvalues, respectively. The dots show the observed pair of values for diffierent contexts (color-coded) and all sources (not annotated). The kernel density estimates represent the distribution for eigenvalues of randomly reshuffled genomic bins. The dashed bisecting line divides the plane into regions where the score is higher in compacted (CSDs) or loose structural domains (LSDs), respectively.

**Figure S6.**
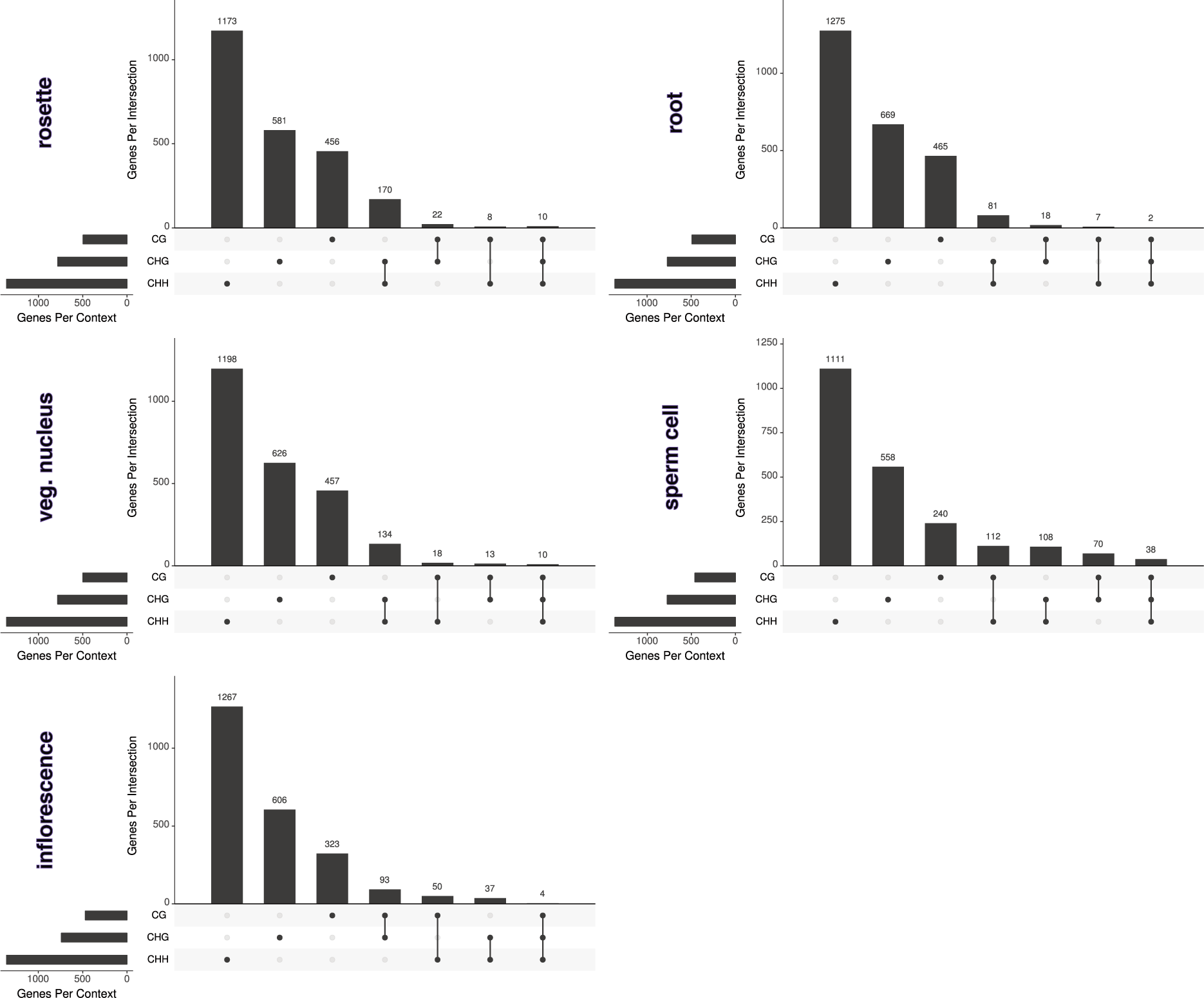
Overlap of top 5% MSGs according by C context. The filled cells below the x-axis indicate the C contexts that are part of the intersection and the y-axis indicates the number of MSGs in the respective intersection. The total number of MSGs in each context is given by the bars on the left, respectively.

**Figure S7.**
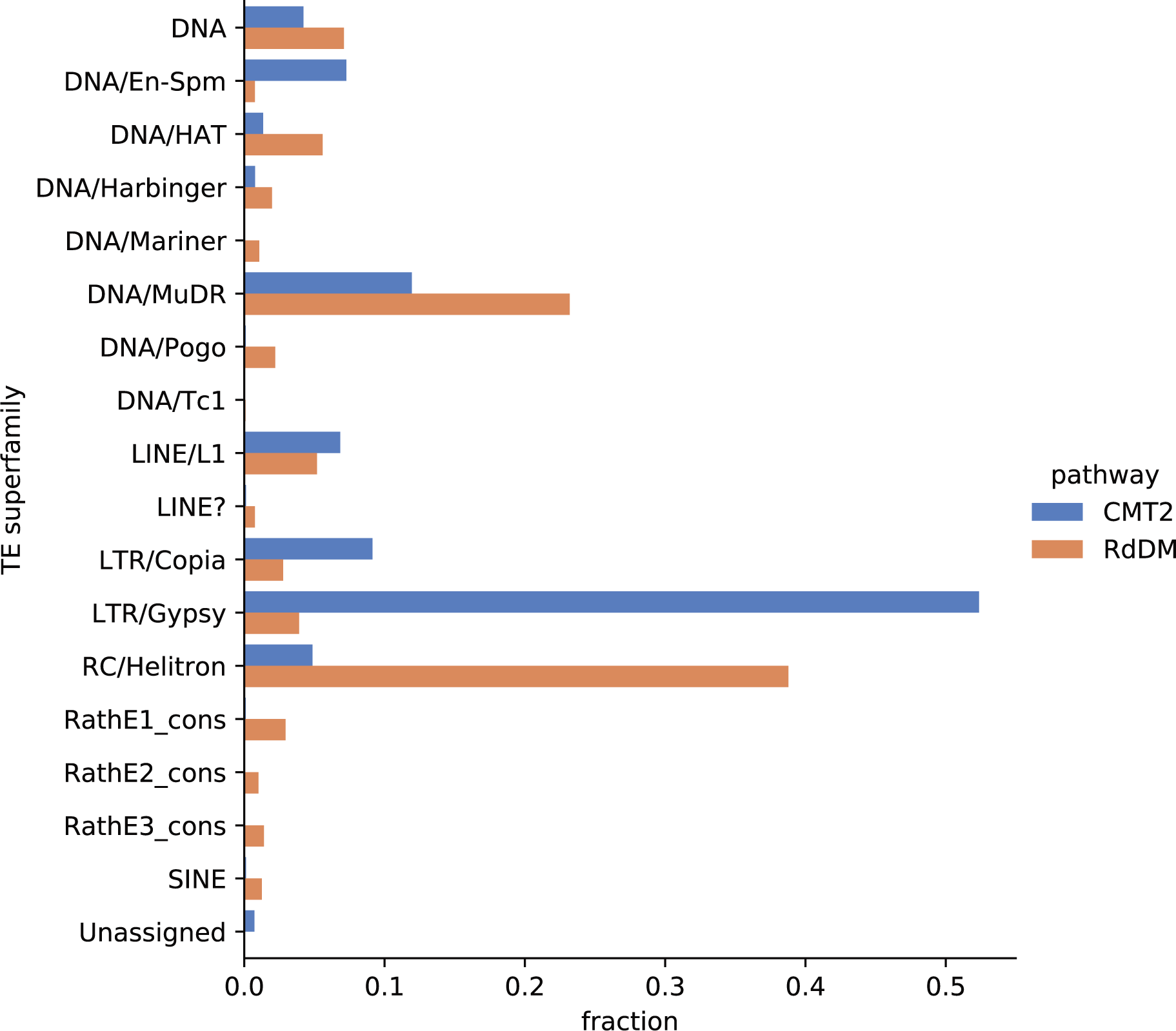
TE superfamily composition of CMT2- and RdDM-targeted TEs. The y-axis lists the TE superfamilies and the x-axis gives the fraction of TEs belonging to a given TE superfamily normalized to the total number of TEs in each DNA methylation category.

**Figure S8.**
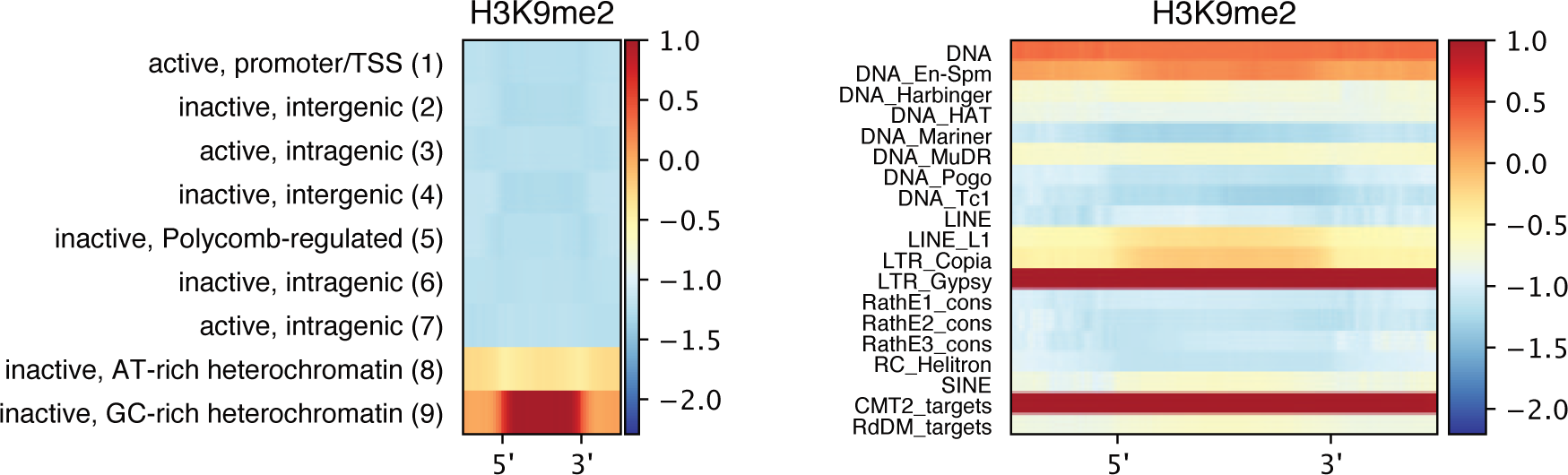
H3K9me2 profiles over chromatin states and TE categories. Each profile comprises 2 kb flanks and color-codes the arithmetic mean of the signal in non-overlapping, 50 bp bins (x-axis). The H3K9me2 signal is the log-transformed average of two replicates normalized to H3, log_2_(H3K9me2/H3), as in [27].

**Figure S9.**
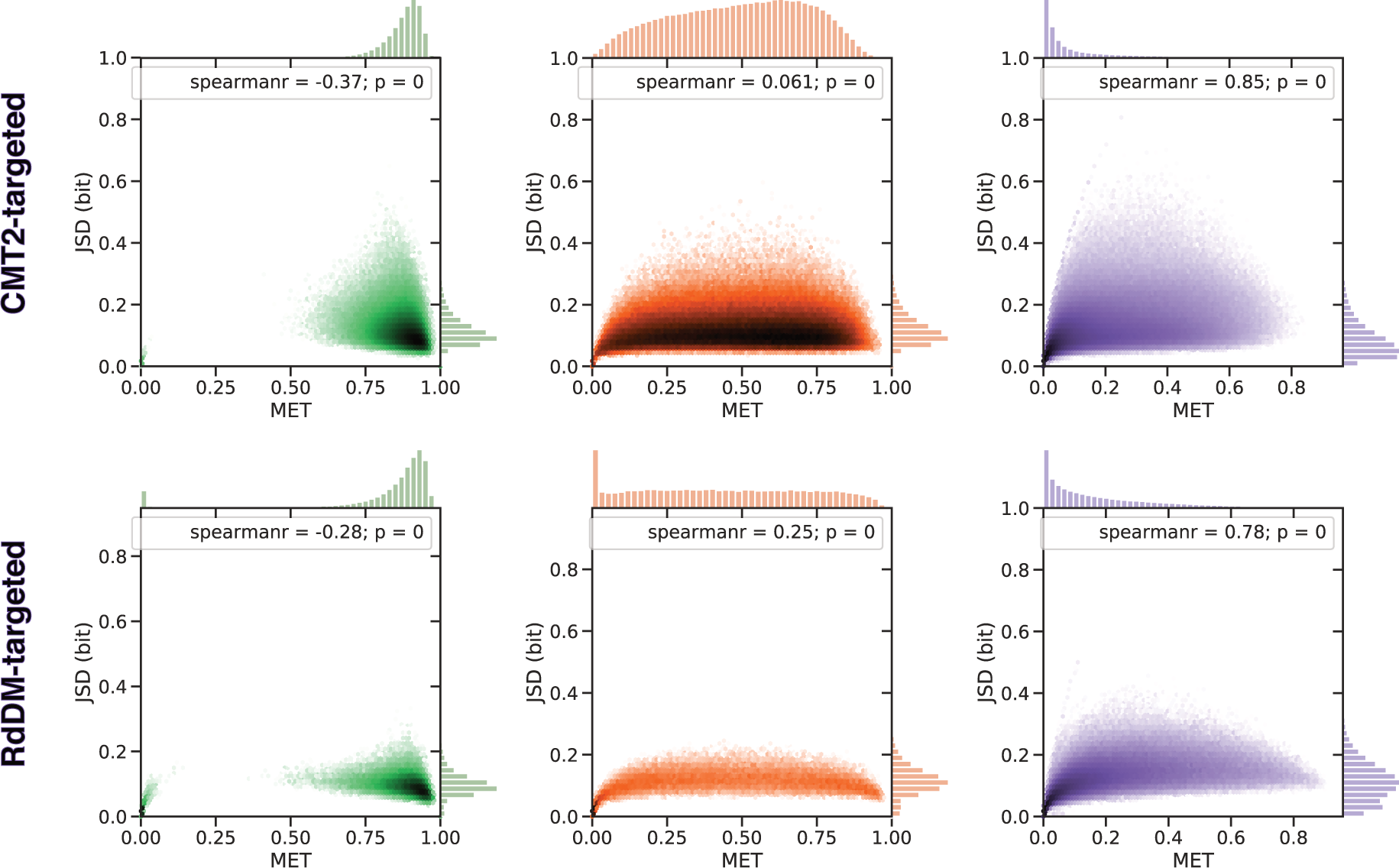
Context-specific MET-JSD phaseplanes for CMT2- and RdDM-targeted TEs. Methylome phase planes for rosette leaves. The margins display the distribution of MET and JSD, respectively. SciPy (scipy.stats.spearmanr) was used to compute the Spearman correlation coefficient and *p*-value.

**Figure S10.**
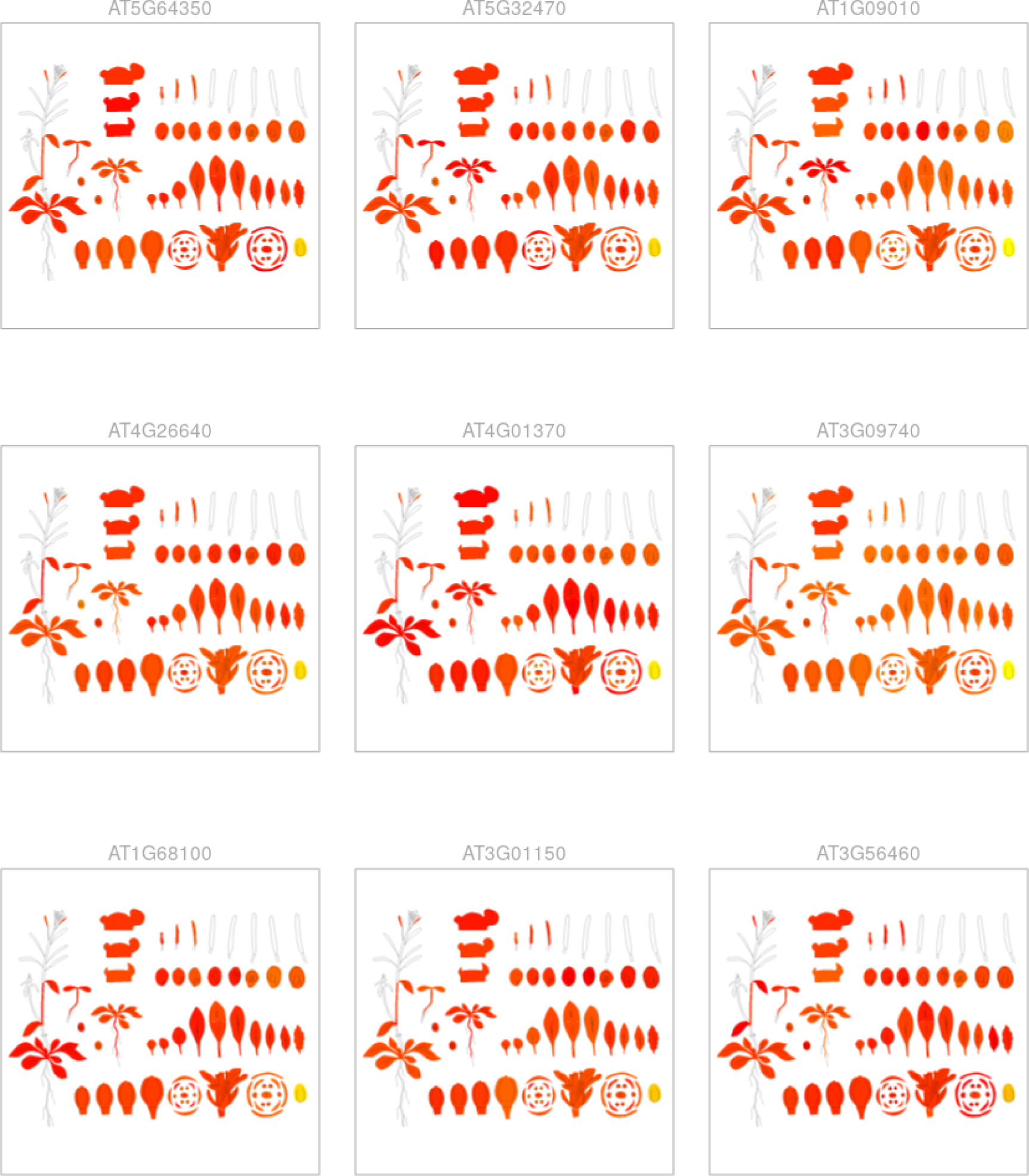
Tissue-specific expression of the bottom 50 genes of mature pollen. The small multiples view of ePlant shows the nine least expressed genes in the bottom 50 gene set of mature pollen. As described on the ePlant website, the gene expression data [82, 83] “generated by the Affymetrix ATH1 array are normalalized [*sic*] by the GCOS method, TGT value of 100.”

